# Multimodal intrinsic activation of GPCRs in ultrastable plasma membrane nanodomains

**DOI:** 10.1101/2024.02.28.582451

**Authors:** Gabriele Kockelkoren, Jens Carstensen, Line Lauritsen, Eleftheria Kazepidou, Asger Tonnesen, Christopher G. Shuttle, Paulina S. Kaas, Ankur Gupta, Artù Breuer, Søren G. F. Rasmussen, Loïc Duffet, Tony Warne, Tommaso Patriarchi, Christopher G. Tate, Mark Uline, Dimitrios Stamou

**Affiliations:** Center for Geometrically Engineered Cellular Membranes, Department of Chemistry, University of Copenhagen, Copenhagen, Denmark; Department of Neuroscience, University of Copenhagen, Copenhagen, Denmark; Institute of Pharmacology and Toxicology, University of Zurich, Zurich, Switzerland; MRC Laboratory of Molecular Biology, Francis Crick Avenue, Cambridge, UK; Department of Chemical Engineering, Biomedical Engineering Program, University of South Carolina, Columbia, South Carolina, United States; Atomos Biotech, Copenhagen, Denmark

## Abstract

G protein-coupled receptors (GPCRs) mediate many physiological functions and are key targets in drug development^1-3^. A long-held tenet of molecular pharmacology is that GPCRs can spontaneously sample preexisting active conformations. This concept is pivotal to our understanding of ligand pharmacology^4^, however, direct evidence supporting it has only been obtained with reconstituted receptors^5-12^. Here, we introduce a method for quantitatively imaging the intrinsic activation probability of GPCRs directly at the plasma membrane of live cells, utilizing fluorescent conformational biosensors^13,14^. Our findings unveil a remarkable spatial multimodality in intrinsic activation probability, with a significant majority (up to 99%) of plasma membrane-expressed receptors showing negligible spontaneous activation. In contrast, the remaining minority of receptors exhibits spontaneous activation up to 22-fold higher than previously estimated. Experiments and theoretical calculations revealed that receptors diffuse into and out of ultralong-lived (∼5 minutes) nanodomains where the local membrane curvature allosterically enhances activation in the absence and presence of ligands. Extensive testing across five prototypic GPCRs indicates spatial nanoscale multimodality is ubiquitous, but varying in magnitude depending on the receptor and cell type. Upending conventional wisdom, this study reveals that drug efficacy is not a constant number but a spatiotemporal function ε *(x, y, z, t)* whose properties define and multiplex the signaling potency and efficacy of ternary complexes of GPCRs and likely other plasma membrane-receptors.

**Graphical Abstract:** 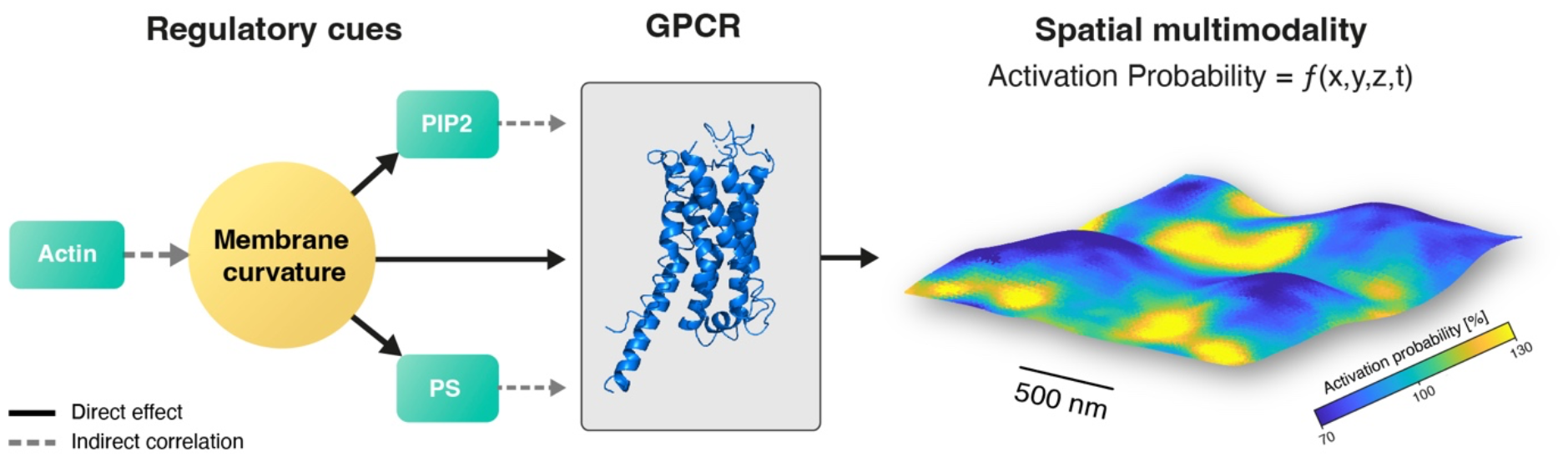

**GPCR spontaneous activation and intrinsic efficacy are not uniform across the plasma membrane but exhibit ultralong-lived spatial multimodality**. Spatial variations in the curvature and composition of the plasma membrane, lead to the emergence of ultralong-lived nanodomains with contrasting physicochemical properties that allosterically regulate GPCR conformations. This results in a multimodal landscape of intrinsic efficacy *ϵ (x, y, z, t)* that ultimately governs cell signaling. XY scalebar: 500 nm. Z-range: 100 nm.

## Introduction

G protein-coupled receptors (GPCRs) represent the largest and most versatile family of plasma membrane receptors and are major pharmaceutical targets^1-3^. A long-held tenet of molecular pharmacology is that GPCRs can interconvert between preexisting active and inactive conformations in the absence of ligands^4^. Ligands are thought to “select” between these conformations, with agonists enhancing the probability of sampling active conformations and inverse agonists reducing it. Thus, the hypothesis of preexisting active conformations underpins our mechanistic understanding of ligand efficacy and pharmacology^4^. However, the direct observations of the basal equilibrium of conformational states that support this hypothesis have been made only in detergent/lipid receptor reconstitutions^5-12^, which do not recapitulate the physiological relevance and complexity of the plasma membrane. Thus, the impact of the plasma membrane on basal conformational equilibria remains unexplored. Here, we developed a method to quantitatively image GPCR intrinsic activation probability (PIA) at the plasma membrane of unperturbed live cells using fluorescent conformational biosensors^13,14^. Our results revealed a pronounced spatial nanoscale multimodality in activation probability, suggesting that in live cells, the distribution of active and inactive conformations is not at equilibrium but under spatial allosteric bias imposed by nanoscale heterogeneities in the curvature and composition of the plasma membrane.

## Results

### Conformational biosensors report intrinsic activation probability (PIA) of β1AR in live cells

As a model receptor for this study, we chose the beta-1 adrenergic receptor (β1AR), a prototypic Gαs-coupled GPCR^15^. To quantify the PIA of β1AR, we used an engineered surrogate of the Gαs subunit (miniG_s_)^16^ which has been widely used to detect the active conformation of β1AR and to study basal and ligand-dependent GPCR activation in live cells with fluorescence microscopy^14,17-19^. MiniG_s_ translocates from the cytoplasm to the plasma membrane, where it reversibly interacts with receptors that sample active conformations (Fig. 1a). At low expression levels, miniG_s_ has been shown to provide a direct proximal readout of GPCR activation that recapitulates ligand efficacy and bias with minimal functional perturbation^14,17-19^.

**Figure 1.**
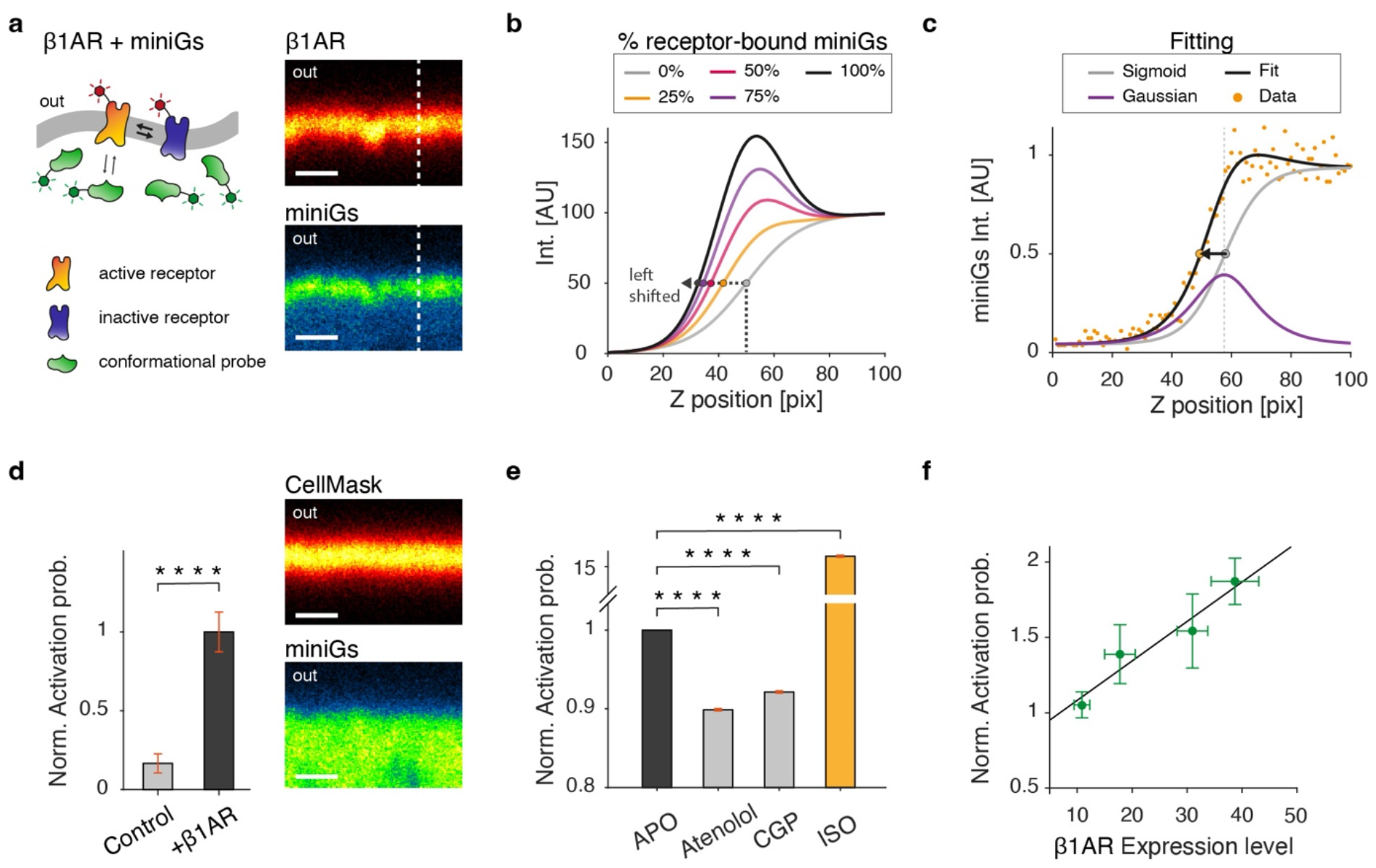
Direct measurement of intrinsic activation probability (P_IA_) with a conformational biosensor. **a**, Left, illustration of interconverting active and inactive receptor conformations at the plasma membrane. A low affinity conformational probe reversibly interacts with the active conformation. Right, confocal XZ line scans of the plasma membrane of HEK-293 cells. Upon activation with full agonist isoproterenol (ISO), SNAP-β1AR (top) transiently recruits GFP-miniG_s_ (bottom). **b**, Simulation of miniG_s_ intensity along a Z line scan of the plasma membrane. The signal can be decomposed into a cytosolic and a receptor-bound contribution respectively represented by a sigmoid and a Gaussian function. Increasing amount of receptor-bound miniG_s_ leads to a left-ward shift of the trace with respect to the Z-position of the membrane. **c**, Typical intensity Z line scan showing the left-ward shift is fitted with the sum of a sigmoid term constraint to the Z-position of the membrane, and a Gaussian term whose amplitude provides a direct measure of miniG_s_ recruitment. **d**, Bar plot of mean miniG_s_ recruitment in cells without and with β1AR expression. Error bars show the s.d.. On the right, confocal XZ line scans of plasma membrane-label CellMask (top) and GFP-miniG_s_ (bottom). miniG_s_ signal is observed in the cytosol. \*\*\*\**P*=6.6×10^-4^, two-sided student *t* test. Cellmask, *n*_*C*_ = 18 and *n*_*R*_ = 3, and for β1AR, *n*_*C*_ = 51 and *n*_*R*_ = 11. **e**, Bar plot showing mean miniG_s_ recruitment to β1AR at the basal state, after addition of inverse agonists atenolol and CGP20712A (CGP), and after activation by ISO. Error bars show the s.e.m. \*\*\*\**P*<10^-10^ (atenolol); \*\*\*\**P*<10^-10^ (CGP) \*\*\*\**P*<10^-10^ (ISO), two-sided student *t* test. For atenolol, *n*_*C*_ = 34 and *n*_*R*_ = 3, for CGP, *n*_*C*_ = 30 and *n*_*R*_ = 3 and for ISO, *n*_*C*_ = 33 and *n*_*R*_ = 4. **f**, Mean β1AR expression level versus mean miniG_s_ recruitment at basal state is plotted into equally spaced bins. Error bars represent the s.e.m and the black line is the linear fit to the data (Pearson’s correlation of 0.96). Each bin contains at least 4 cells, and *n*_*C*_ = 22 and *n*_*R*_ = 11. Scalebar for all micrographs, 1 μm. *P*>0.05 is not significant (n.s.), while *P*<0.05 is significant. Hereafter, *n*_*C*_ represents the number of cells, and *n*_*R*_ represents the number of biological replicates. All images herein were acquired in standard confocal mode unless otherwise stated.

To measure quantitatively P_IA,_ we simultaneously imaged miniG_s_ and β1AR labeled with two spectrally distinct fluorophores using confocal microscopy (Fig. 1a). We then measured axial intensity profiles of the receptor and the miniG_s_ channel (Fig. 1a, b). Axial profiles of the receptor channel are fitted with a Gaussian function whose amplitude provides relative receptor density. Axial profiles of miniG_s_ can be described by the sum of a Gaussian and a sigmoid, which respectively represent the relative receptor-bound and cytosolic fraction of miniG_s_ (Fig. 1b, c). To measure the Gaussian term with greater sensitivity, we leveraged the precise localization of the axial position of the plasma membrane in the receptor channel (± 3 nm)^20^ which allowed us to fix the Z-position of the Gaussian term (Fig. 1c, Extended Data Figs. 1 and 2, see detailed description in Methods and Supplementary information). This automated analysis provided the background-corrected relative density of miniG_s_ and receptor. Their ratio along the plasma membrane is a direct measure of relative P_IA_.

**Figure 2.**
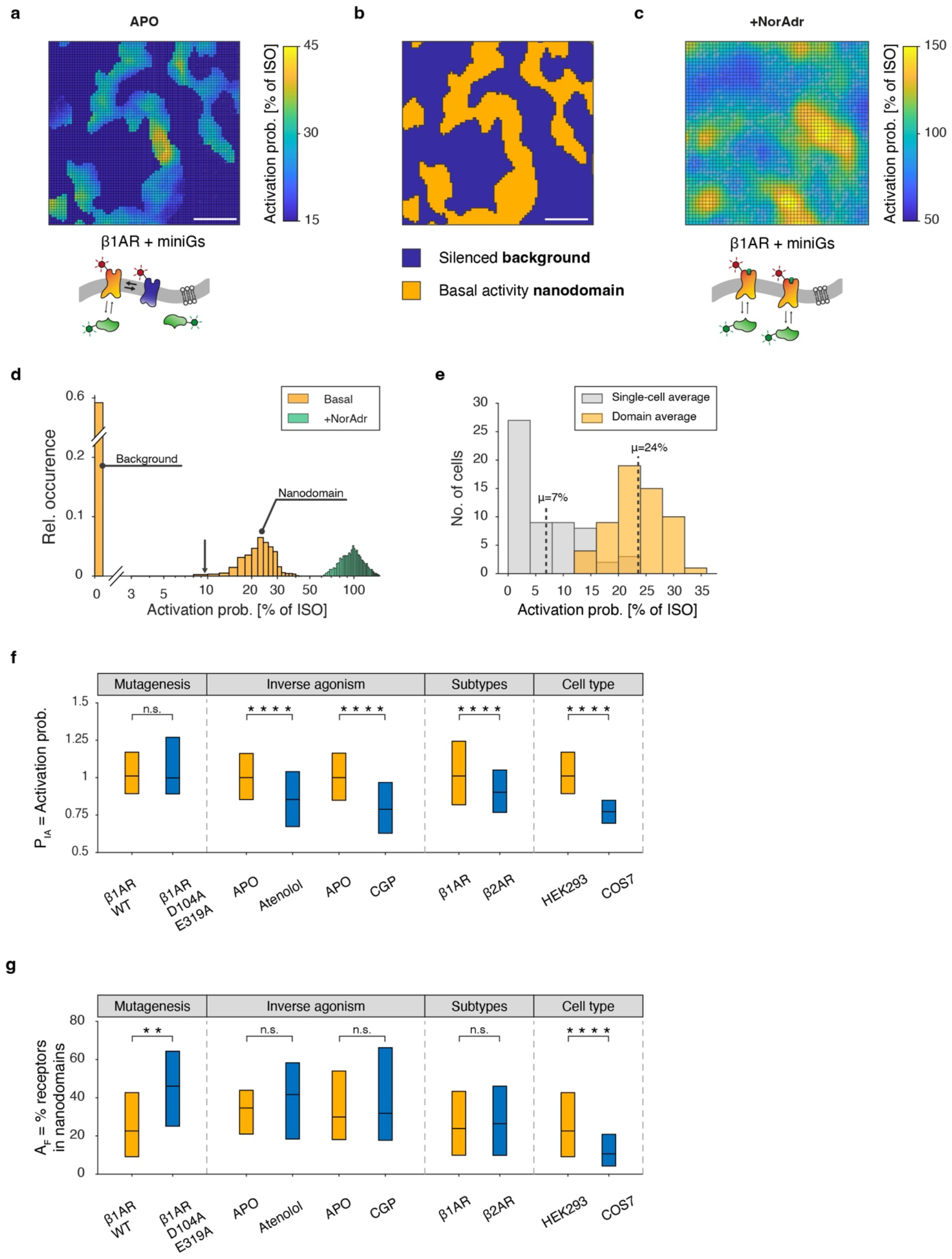
Plasma membrane nanodomains of multimodal β1AR spontaneous activation and intrinsic efficacy. **a**, Typical spatially resolved map of probability of spontaneous/basal intrinsic activation (P_IA_) of β1AR at the adherent plasma membrane of HEK293. **b**, Thresholding of (a) allows quantification of background areas (blue) where spontaneous P_IA_ ∼ 0 and areas of nanodomains (orange) where spontaneous P_IA_ ∼ 24%. **c**, Typical map of β1AR spontaneous P_IA_ after addition of full agonist noradrenaline. Spatial multimodality persists even though all receptors are now activated. Color-scale (a, c) is normalized to the average activation induced by ISO. Illustrations of the experimental setup below the maps in (a, c). Scale bar (a, c), 500 nm. **d**, Histograms of spatially resolved P_IA_. Whole-cell average of P_IA_ is indicated by black arrow. **e**, Histograms of average P_IA_ for entire cells (i.e., domains and background) (grey) and for domains only (yellow). Dashed lines indicate means of each population. Bulk assays read out an average P_IA_ of 6.9% and, thus, assume each receptor has a 6.9% probability to activate. On average, basal active nanodomains have a probability of activation of 23.6%, whereas receptors in the background ∼0%. Data in (d), *n*_*C*_ = 58 and *N*_*R*_ = 10. **F, g**, Boxplots for P_IA_ (f) and A_F_ (g) show differences between β1AR WT and a double mutant, between basal (APO) state and addition of inverse agonists atenolol and CGP12712A, between β1AR and β2AR, and between basal β1AR state in HEK293 and COS7 cells. For P_IA,_ n.s. *P*=0.62 (mutagenesis); \*\*\*\**P*<10^-10^ (atenolol); \*\*\*\**P*<10^-10^ (CGP); \*\*\*\**P*<10^-10^ (subtypes); \*\*\*\**P*=1.7×10^-10^ (cell types). For A_F_, \*\**P*=4.0×10^-3^ (mutagenesis); n.s. *P*=0.17 (atenolol); \*\*\*\**P*=0.58 (CGP); n.s. *P*=0.84 (subtypes); \*\*\*\**P*=9.0×10^-4^ (cell types). Significance test is two-sided Kolmogorov-Smirnov. Data is β1AR and miniG_s_, *n*_*C*_ = 82 and *n*_*R*_ = 11. For β1AR APO in HEK293 cells, *n*_*C*_ = 82 and *n*_*R*_ = 11, for β1AR D104A/E319A, *n*_*C*_ = 39 and *n*_*R*_ = 4, for β1AR APO and atenolol, *n*_*C*_ = 34 and *n*_*R*_ = 3, for β1AR APO and CGP, *n*_*C*_ = 30 and *n*_*R*_ = 3, for β2AR APO, *n*_*C*_ = 65 and *n*_*R*_ = 9, for β1AR APO in COS7 cells, *n*_*C*_ = 51 and *n*_*R*_ = 4.

To validate the method, we performed several negative and positive controls. Expression of β1AR under apo conditions increased the plasma membrane recruitment of miniG_s_ by 600% (Fig. 1d), confirming the specific detection of the very small basal/spontaneous P_IA_ and revealing that our detection limit is 6-fold smaller than basal activation. This is a 20-to 60-fold improvement in sensitivity compared to ensemble-average methods^21,22^. Consistent with cAMP-based measurements of β1AR activation^23,24^, atenolol and CGP-20712A (CGP) reduced basal P_IA_ by ∼10%, while addition of the full agonist isoproterenol (ISO) increased basal P_IA_ by ∼1500% (Fig. 1e). Finally, miniG_s_ basal recruitment was measured as a function of receptor expression level and showed the expected linear increase^14,22^ (Fig. 1f). Collectively, these results demonstrate the direct, specific and highly sensitive detection of relative P_IA_ for β1AR in the presence and absence of agonist.

### Multimodal β1AR intrinsic activation in plasma membrane nanodomains

To generate high-resolution 3D maps of P_IA_ at the plasma membrane, we combined XZY confocal scans as described in detail in reference 20. Briefly, we used axial localization microscopy to reconstruct with ∼3 nm axial precision the 3D surface of the plasma membrane^20^. We calculated spatial maps of P_IA_ at the plasma membrane as the stoichiometry of miniG_s_:β1AR, thus accounting for heterogeneities in receptor density^20,25^. To quantify P_IA_, we normalized it to the maximal recruitment of miniG_s_ induced by the full agonist isoproterenol (ISO) on per single-cell basis (see Methods and Supplementary Methods).

The current model of basal signaling posits that spontaneous receptor activation is a stochastic process driven by thermal energy^4^. Based on this stochastic model, one would hypothesize that P_IA_ is relatively uniform across the plasma membrane and exhibits a Gaussian distribution around a mean value (Extended Data Figs. 3a and b). Intriguingly, however, our data reveal a striking spatially multimodal distribution of P_IA_. We observe a highly heterogenous map consisting of clearly defined nanodomains of high P_IA_ surrounded by ‘background’ areas of high receptor density but where P_IA_ was below the detection limit of 1.2% (Figs. 2a, b and Extended Data Fig. 3c). The evident multimodality of the images can be quantified in histograms of P_IA_ that reveal two major, completely separated, populations centered around ∼0% and ∼24% (Fig. 2d, orange).

**Figure 3.**
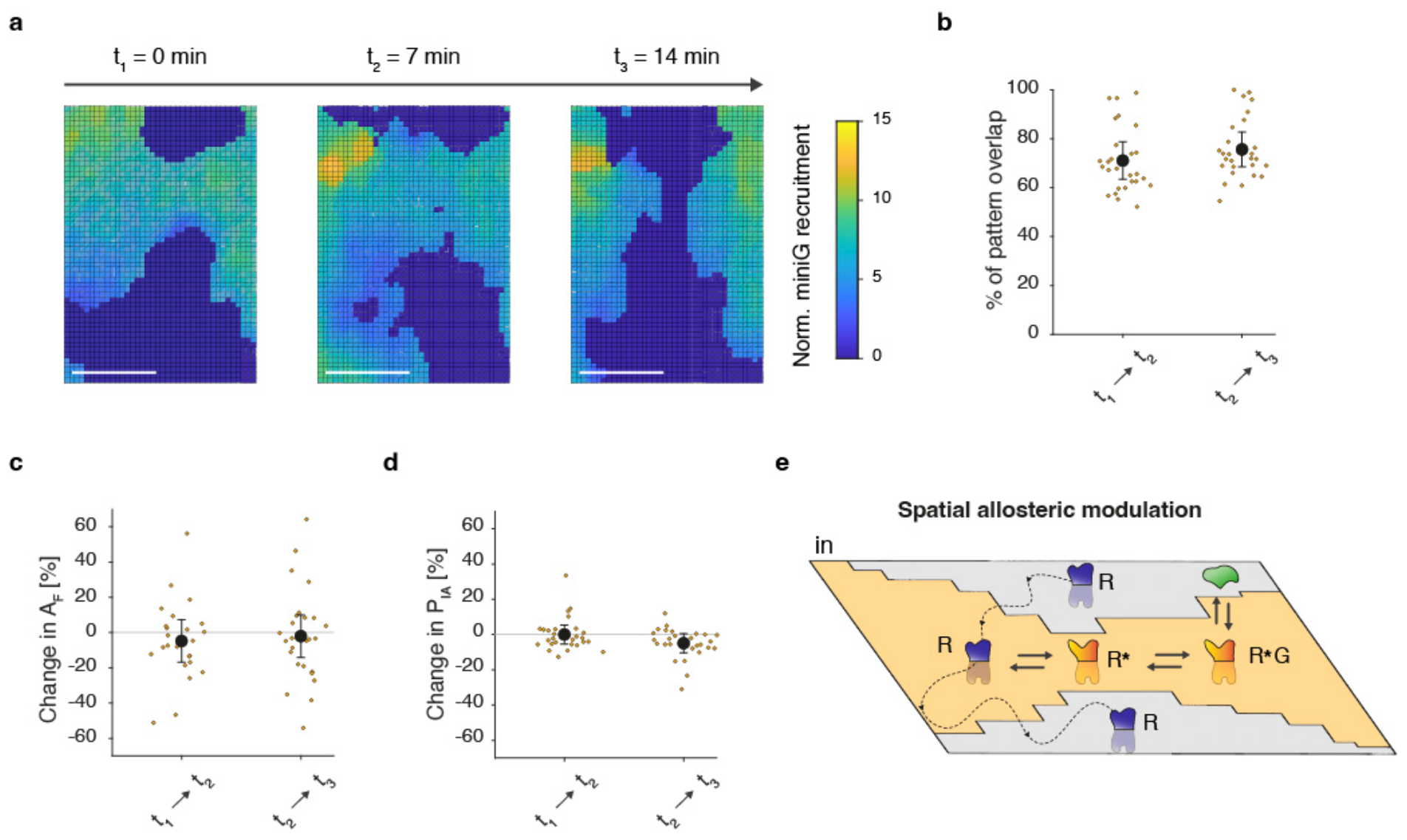
Ultralong-lived spatial multimodality of β1AR intrinsic activation probability. **a**, Consecutive maps of P_IA_ at the same region of the plasma membrane. Data is representative of n_c_ = 27 and n_R_ = 3. **b, c, d**, Percentage change between consecutive maps, respectively, for percentage of overlapping pixels (b), A_F_ (c) and P_IA_ (d). P_IA_ patterns exhibit ultralong-lived spatial stability (∼75% over ∼5min). Data is shown as mean +/-s.e.m. and yellow points represent single cells. Data is n_c_ = 27 and n_R_ = 3. **e**, Schematic illustrating how the plasma membrane acts as a spatial allosteric modulator for inactive (R) and active (R*) conformational states. In the background (grey area) receptors are ‘locked’ in the inactive R state. Fast diffusing receptors from the background into nanodomains (orange area) interconvert to the active state. In nanodomains, receptors in the active state R* can subsequently bind to a G protein. Diffusion from a nanodomain into the background interconverts the receptor back into the inactive state.

We validated the multimodal distribution of P_IA_ reported by miniG_s_ with two additional sensors. First, we employed a widely-used conformation-specific biosensor for activated β1AR known as nanobody 80 (Nb80)^13^ which quantitatively confirmed the miniG_s_ findings (c.f. Fig. 2d, e and Extended Data Fig. 4). Second, we employed a conformational biosensor for dopamine receptor D1 (DRD1) based on circularly permutated GFP^26^ which recapitulated pronounced spatial multimodality in P_IA_ for DRD1 (Extended Data Fig. 5). The latter control excludes the possibility that multimodality is due to restricted accessibility of receptors to cytosolic biosensors or due to miniG_s_-impaired internalization^27^. We also performed ratiometric imaging of two spectrally distinct membrane dyes which revealed a uniform spatial distribution confirming the absence of putative imaging artifacts in P_IA_ (Extended Data Fig. 11).

**Figure 4.**
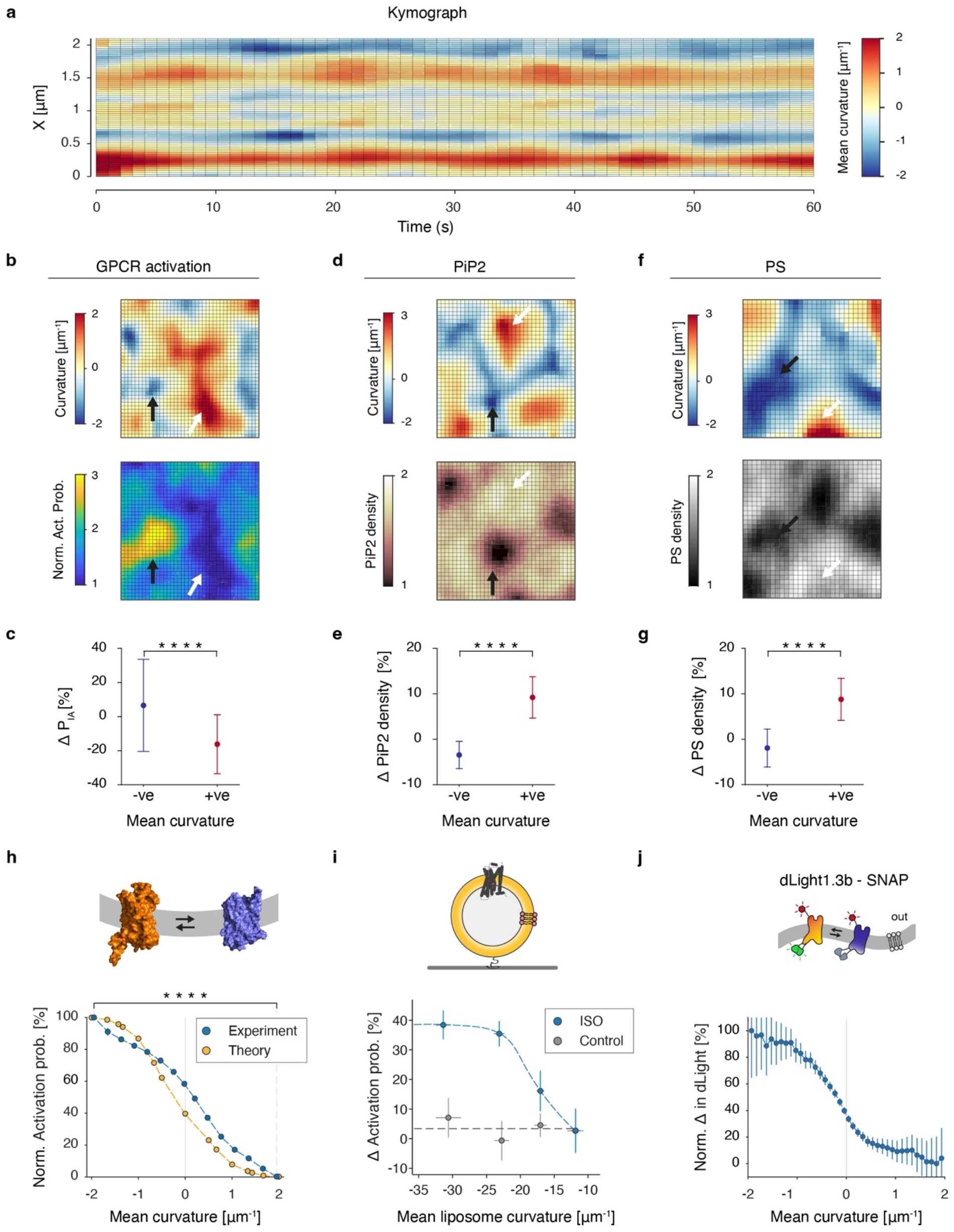
Membrane curvature regulates receptor activation probability. **a**, To assess changes in membrane curvature over time, we produced one dimensional kymographs (x-axis) of membrane curvature over time. Typical kymograph over t = 60 s. Within this time frame, most curvature features are stable. **b-g**, Map of mean curvature and β1AR receptor activation probability (b), PIP2 density (d) and PS density (f). Arrows point at representative regions of colocalization with positive and negative curvature (white and black arrow, respectively). Change in β1AR receptor activation probability (c), PIP2 density (e) and PS density (g) between negative and positive curvature. Data is mean +/-SD for single cells. \*\*\*\**P*=7.5×10^-9^ (P_IA_); \*\*\*\**P*=4.0×10^-16^ (PIP2); \*\*\*\**P*=3.9×10^-18^ (PS), two-sided student *t* test. For β1AR, *n*_*C*_ = 82, *n*_*R*_ = 11; for PIP2, *n*_*C*_ = 27, *n*_*R*_ = 3; for PS, *n*_*C*_ = 44, *n*_*R*_ = 3. **h**, Mean curvature versus β1AR receptor activation probability measured at the plasma membrane of live cells (blue) and by theoretical FT calculations (yellow). Data are binned using an error-weighted rolling average (0.3 ± 0.3 μm^-1^) with error bars shown as s.e.m. \*\*\*\**P*<10^-10^, two-sided student *t* test. For β1AR, *n*_*C*_ = 44, *n*_*R*_ = 7. **i**, Change in activation probability, measured as changes in BDY intensity, as a function of liposome membrane curvature for control with buffer (grey data) and upon addition of ISO (blue data). Each bin represents mean +/-s.e.m. and contains 1000–1300 liposomes. For ISO, *n*_*R*_ = 8 and for control *n*_*R*_ = 3. **j**, Normalized change in of dLight1.3b (measured as ratio of cpGFP / SNAP) as a function of mean membrane curvature. Measurements were performed at saturating concentrations of dopamine (10 μM). Data are binned using an error-weighted rolling average (0.1 ± 0.1 μm^-1^) with error bars showing s.e.m. Data is representative of *n*_*C*_ = 55, *n*_*R*_ = 4.

**Figure 5.**
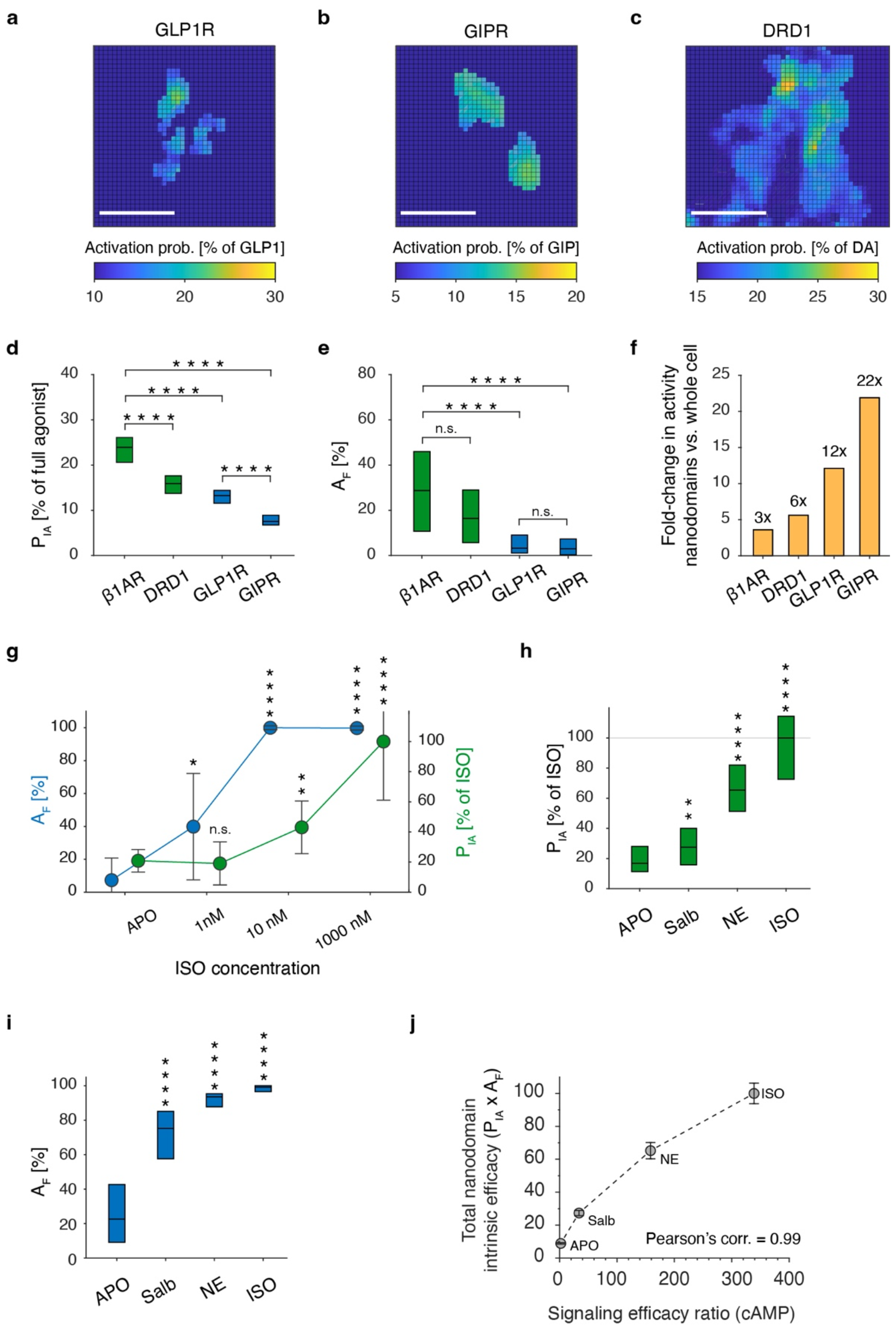
Spatial functional multimodality is a general, ligand-dependent, and titratable property of GPCRs. **a, b, c**, Maps of multimodal activation probability normalized to the response induced by canonical full agonists GLP1, GIP and dopamine (DA). Scale bar, 500 nm. **d, e**, Boxplots for P_IA_ (d) and A_F_ (e) show differential regulation of P_IA_ and A_F_ between receptors. *P* values are calculated by two-sided Kolmogorov-Smirnov test. *P*>0.05 is not significant (n.s.), while *P*<0.0001 is significant (4 stars). For P_IA,_ \*\*\*\**P*=2.3×10^-10^ (β1AR-DRD1); \*\*\*\**P*=2.4×10^-14^ (β1AR-GLP1R); \*\*\*\**P*=2.9×10^-15^ (β1AR-GIPR); \*\*\*\**P*=3.0×10^-8^ (GLP1R-GIPR). For A_F,_ n.s. *P*=0.11 (β1AR-DRD1); \*\*\*\**P*=1.5×10^-6^ (β1AR-GLP1R); \*\*\*\**P*=7.8×10^-9^ (β1AR-GIPR); n.s. *P*=0.13 (GLP1R-GIPR). **f**, Fold-difference in GPCR basal activity in nanodomains compared to whole-cell averages for 4 different receptors. **g**, Titration of nanodomain properties P_IA_ and A_F_ by agonist isoproterenol (ISO). Data is mean +/-sd. **h, i**, Boxplots for P_IA_ (g) and A_F_ (h) show how nanodomains differentially regulate active and inactive receptors in a ligand-dependent manner for salbutamol (Salb), norepinephrine (NE) and ISO. All ligands have been added at saturating concentrations and P_IA_ is normalized to ISO. **j**, Total nanodomains intrinsic efficacy scales with cAMP signaling efficacy ratio (from ref. ^47^). APO efficacy ratio is approximated as basal cAMP signaling levels normalized to ISO. Data is mean +/-sem. For P_IA,_ \*\*\*\**P*=0.075 (APO-Salb); \*\*\*\**P*=1.3×10^-8^ (APO-NE); \*\*\*\**P*=2.6×10^-10^ (APO-ISO). For A_F,_ \*\*\*\**P*=1.0×10^-10^ (APO-Salb); \*\*\*\**P*=1.2×10^-10^ (APO-NE); \*\*\*\**P*=5.0×10^-13^ (APO-ISO). **i**, Titration of nanodomain properties P_IA_ (green) and A_F_ (blue) with ISO. Data points represent the mean over all cells +/-sd. For P_IA,_ \*\*\*\**P*=0.9575 (APO-1nM); \*\*\*\**P*=0.0035 (APO-10nM); \*\*\*\**P*=5.4×10^-5^ (APO-1000nM). For A_F,_ \*\*\*\**P*=0.019 (APO-1nM); \*\*\*\**P*=5.4×10^-5^ (APO-10nM); \*\*\*\**P*=5.4×10^-5^ (APO-1000nM). Data β1AR and miniG_s_, *n*_*C*_ = 58 and *n*_*R*_ = 10, GLP1R and miniG_s_, *n*_*C*_ = 33 and *n*_*R*_ = 3, GIPR and miniG_s_, *n*_*C*_ = 28 and *n*_*R*_ = 3, and DRD1 and miniG_s_, *n*_*C*_ = 22 and *n*_*R*_ = 3. Data APO, *n*_*C*_ = 82 and *n*_*R*_ = 11, Salb, *n*_*C*_ = 38 and *n*_*R*_ = 3, NE, *n*_*C*_ = 19 and *n*_*R*_ = 3, and ISO, *n*_*C*_ = 16 and *n*_*R*_ = 3. Data ISO titration on β1AR and miniG_s_, *n*_*C*_ = 9 and *n*_*R*_ = 2.

Adding the full agonist noradrenaline at saturating concentration conjoins the background-with the nanodomain-population into a single distribution centered at P_IA_∼100% (Fig. 2d, green). This crucial control demonstrates that receptors in background areas are not in some way ‘defective’ but can be activated by agonists. It also excludes the possibility that multimodality is due to the depletion of miniG_s_ or restricted accessibility of the receptor to miniG_s_. Intriguingly, this control also demonstrates that even though all receptors activate at saturating agonist concentration, P_IA_ does not become spatially uniform (Fig. 2c). We confirmed this observation for β1AR expression levels varying over almost 2 orders of magnitude (Extended Data Fig. 12) and at the freestanding (i.e. apical) membrane of cells (Extended Data Fig. 13). Thus, multimodality is also a property of the ligated receptor and, thus of intrinsic efficacy.

Overall, our findings suggest that whole-cell measurements of receptor activation conceal a multimodal spatial distribution of receptors with distinct intrinsic activation. Such a whole-cell average is proportional to the product A_F_ × P_IA_, where A_F_ is the fraction of activated receptors (i.e., receptors in nanodomains) and P_IA_ is their activation probability (Fig. 2e).

Next, we performed controls well-known to increase/decrease spontaneous receptor activity in whole-cell experiments to deconvolute their dependence on spatial multimodality. To increase spontaneous activation, we used a β1AR double mutant with high spontaneous activity (D104A^2.50^, E319A^6.30^)^22,28^, while to reduce it we used the inverse agonists CGP and atenolol^23,24^ (Fig. 2g, h). Both controls increased/decreased, respectively, whole-cell activation probability as expected. Intriguingly, however, they exploited distinct mechanisms. While the inverse agonists regulated solely P_IA_, the double mutation regulated solely A_F_, thus demonstrating these two distinct metrics of spatial multimodality are independent regulatory cues of spontaneous (or constitutive) activity (Fig. 2g, h).

Spontaneous activity has also been broadly reported to depend on receptor subtype and cell type. As a further test, we thus compared the P_IA_ of β1AR versus β2AR in HEK293 and β1AR in HEK293 cells versus COS7 cells (Fig. 2g, h). These quantitative comparisons were made possible because we leveraged ratiometric β1AR/miniG_s_ measurements to quantify the probability of receptor activation. This metric, by definition, does not depend on receptor expression level, in contrast to the readouts of most classical signaling assays^22^. These experiments revealed that β1AR has ∼ 10% higher P_IA_ than β2AR but statistically indistinguishable A_F_. It is worth mentioning that though there is agreement in the literature that the two subtypes exhibit different spontaneous signaling, the relative levels of spontaneous activity are irreproducible^29,30^. β1AR exhibited both higher P_IA_ and AF in HEK293 cells compared to COS7 cells (respectively +24% and +12%, Fig. 2g, h). Historically, variations in signaling between cell types are attributed to changes in the expression/stoichiometry of signaling partners^30^, however, our experiments identify multimodality in intrinsic activation as a causal underlying effector.

Collectively, the exhaustive control experiments in this section validate the existence of multimodal intrinsic activation, thus revealing an intricate network of hitherto unsuspected spatial regulation that manifests through the plasma membrane and thus, by definition, does not exist and cannot be observed in GPCRs extracted out of cellular membranes.

### β1AR interconverts between active/inactive conformations while diffusing in/out of nanodomains

To investigate whether the spatial patterns of P_IA_ arise from confined/trapped biochemically distinct receptor species, we compared the temporal stability of spatial multimodality to the kinetics of receptor diffusion. Consecutive maps of P_IA_ at the same region of the plasma membrane revealed multimodality patterns were mostly conserved over periods of ∼5 min (Fig. 3a and Extended Data Fig. 6c). Indeed, nanoscale P_IA_ patterns recorded 7 min apart had an astonishing ∼75% spatial overlap and no detectable change in the average A_F_ and P_IA_ (Fig. 3b-d), thus demonstrating ultralong-lived temporal conservation. In stark contrast, fluorescence recovery after photobleaching experiments showed that throughout ∼7 min, receptors were exchanged and intermixed ∼400 times over because of their very fast diffusion coefficient^25,31,32^ (∼ 0.06 μm^2^/s, see Extended Data Fig. 6 and Methods). Consequently, the spatial patterns of P_IA_ cannot arise from trapped putative biochemical receptor species of contrasting P_IA_. Our data suggest a different model where the conformational equilibrium of fast diffusing receptors is allosterically biased as they randomly enter and exit ultralong-lived nanodomains. This would be consistent with the μs-ms conformational interconversion rates measured by NMR^5,7^ and single molecule experiments^9,33^.

**Figure 6.**
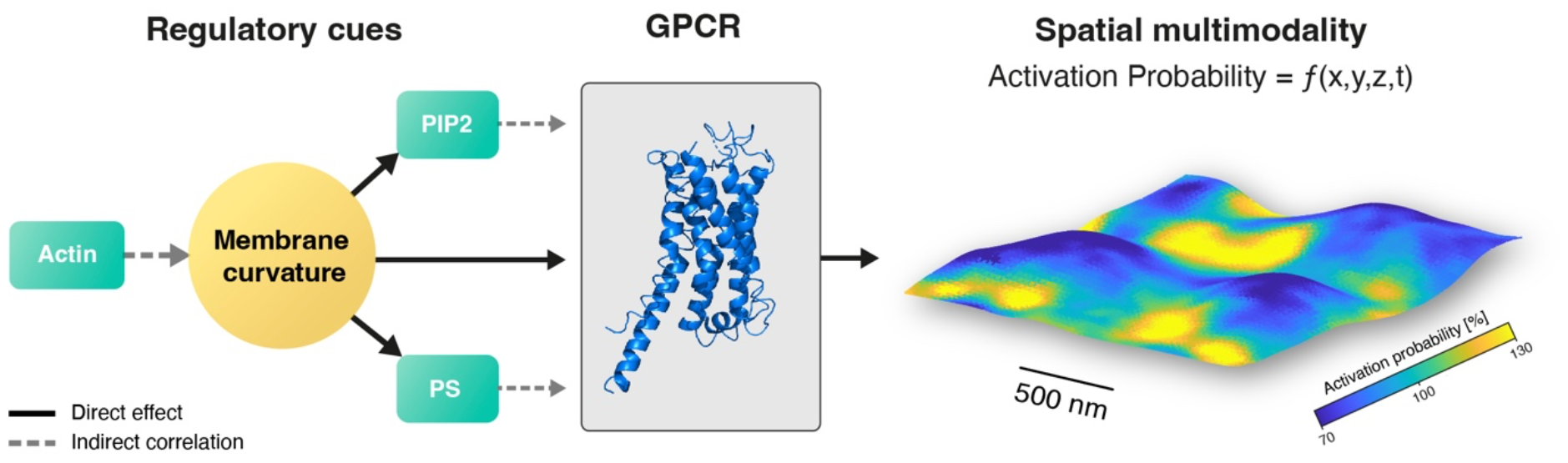
Illustration of the regulatory cues that underpin GPCR spatial functional multimodality. Nanodomains of activated GPCRs are primarily regulated by the curvature of the plasma membrane, potentially in synergy with actin and the anionic lipids PIP2 and PS. The heterogeneous spatial distribution of these regulatory cues creates a three-dimensional multimodal landscape of intrinsic activation probability at the plasma membrane *ε (x, y, z, t)* that ultimately governs cell signaling. Scalebar, 500 nm in XY and 100 nm in Z.

### Membrane curvature allosterically modulates receptor activation probability

To illuminate the molecular mechanism that could enable ultralong-lived nanoscale patterns of allosteric modulation, we hypothesized here that membrane curvature-mediated biophysical interactions could allosterically modulate the conformational equilibria of GPCRs like others have reported, e.g., for bilayer thickness^34,35^. This hypothesis was motivated by recent findings that the spatial organization of GPCRs^20^ is overwhelmingly determined by subtle undulations in the shape/curvature of the plasma membrane.

Kinetic measurements of plasma membrane topography (see Methods and ^20^) confirmed the existence of ultralong-lived nanoscale curved domains, which maintained their overall curvature for periods up to ∼10 min (Fig. 4a and Extended Data Fig. 7). Consistent with the role of the cytoskeleton in maintaining cellular morphology^36^, we found membrane curvature correlated partially with actin density (Extended Data Fig. 8)^20^, which also remained globally stable on these timescales (Extended Data Fig. 14). Notably, in agreement with our hypothesis, we discovered a quantitative correlation of P_IA_ with mean curvature, where β1AR activation scaled with increasing negative curvature (Fig. 4b, black arrow, and 4c). However, additional experiments revealed that plasma membrane curvature correlates also with the spatial organization of negatively charged lipids known to affect receptor function^37-39^ (phosphatidylinositol 4,5-bisphosphate (PIP2) and phosphatidylserine (PS), Figs. 4d-g). To deconvolute the putative synergistic contribution of nanodomain curvature and lipid composition to GPCR activation probability, we leveraged molecular field theory (MFT)^40,41^.

MFT has been shown to provide accurate quantitative predictions of curvature sensing by membrane binding proteins^42-44^ and GPCRs^20^. Here, we expanded the MFT model to include atomistic details of the inactive and active β1AR structures. Thus, we calculated the relative equilibrium probability of β1AR as a function of mean curvature (Fig. 4h, yellow, and Supplementary Methods). The model predicted a monotonic, S-shaped, inverse dependence of P_IA_ on mean curvature. To understand the molecular origin of this behavior, we mapped changes in receptor density and activation probability by membrane curvature onto the structure of β1AR (Extended Data Fig. 15). We find that allosteric activation by curvature relies on the differential perturbation of specific amino acids, which, crucially, have been previously identified to affect receptor activation through ensemble-average mutagenesis studies (Supplementary Methods). Taken together, our results suggest that negative curvatures are sufficient in silico to allosterically enhance P_IA_.

To test and quantitatively compare this theoretical prediction with live-cell experiments, we superposed the spatial patterns of P_IA_ with the underlying mean curvature of the plasma membrane (as shown in Fig. 4b) for >600,000 random locations and performed a correlation analysis. In astonishing agreement with the theory, all experimental data points collapse into an inverted S-shaped curve (Fig. 4h, blue). This analysis confirms with high statistical significance (P<10^-10^) that progressively negative curvatures are necessary to enhance P_IA_ at the plasma membrane. Overall, the remarkable quantitative agreement between experimental and theoretical correlation curves (Fig. 4h) suggests that the mean curvature of the plasma membrane is necessary and sufficient to regulate the P_IA_ of β1AR.

To validate further the finding that membrane curvature regulates receptor P_IA,_ we performed additional control experiments using two entirely different reporters of activation probability. First, we used the well-established model system of β2AR, reconstituted into lipid vesicles and labeled at residue Cys265 with an environmentally sensitive fluorophore (Methods)^45,46^. Indeed, the activation of reconstituted β2AR increased with negative membrane curvature, in agreement with the tendency observed in the plasma membrane (albeit for ∼10-fold smaller mean curvatures, Fig. 4j, Extended Data Fig. 9). Second, we leveraged the well-tested circularly permuted (cp) GFP sensor of DRD1 activation^26^, which confirmed a curvature dependence of P_IA_ in quantitative agreement with miniG_s_ (Fig. 4i, Extended Data Fig. 5). Since these two experiments were done at saturating conditions of full agonist (respectively isoproterenol and dopamine) they also confirm that the curvature-imposed spatial allosteric bias persists for ligand-bound receptors.

### Spatial multimodality as a general but specific mechanism that underpins GPCR potency and efficacy

The intimate energetic coupling of membrane curvature and protein structure, led us to hypothesize that the spatial features of the nanodomains may adapt to differences/changes of receptor sequence/structure. This is a prediction that we tested with three different types of experiments.

First, we investigated whether spatial multimodality is present in other classes of GPCRs and whether it exhibits sequence/structure specificity. We thus measured two prototypical class B receptors, the glucagon-like peptide 1 receptor^48^ (GLP1R) and the gastric inhibitory polypeptide receptor (GIPR). The results convincingly demonstrated spatially heterogeneous P_IA_ for both GLP1R and GIPR (Figs. 5a and b). Furthermore, as expected, the two class B receptors had significantly lower intrinsic activation than the two class A receptors we studied here (β1AR and DRD1). Intriguingly, each receptor exhibited quantitively distinct spatial multimodality features, differing both in the percent of receptors that were spontaneously/constitutively activated (A_F_) and in the intrinsic activation probability they exhibited (P_IA_) (Figs. 5d, e). It is noteworthy that when receptors are activated, they exhibit an intrinsic basal activation probability that is 3-fold to 22-fold higher than a whole-cell average measurement would suggest (Fig. 5f). Taken together, these results indicate that spatial multimodality is a general, yet receptor-specific, phenotype of functional organization for GPCRs.

Second, we measured multimodality at increasing concentrations of β1AR agonist ISO and showed that nanodomain properties are titratable (Fig. 5g), with increasing concentrations of ISO resulting in a monotonic increase in P_IA_ and A_F_. These results represent important positive controls that validate our ability to measure concentration-dependent differences in nanodomains. Furthermore, they reveal that P_IA_ and A_F_ can be regulated independently from each other. Finally, and most crucially, the ligand-dependent evolution of nanodomain properties demonstrates that spatial multimodality underlies classical, ensemble-average, dose-response curves and thus measurements of drug potency.

Third, to establish the regulatory role of spatial multimodality in GPCR activation, we studied three well-known adrenergic ligands. Our results reveal significant and ligand-specific differences in intrinsic activation probability, P_IA_ (Fig. 5h,), and the percentage of receptors in nanodomains, A_F_ (Fig. 5j). For example, while salbutamol exerts its action predominantly via A_F_, ISO leverages both P_IA_ and A_F_. These results suggest that A_F_ and P_IA_ encode drug efficacy in an agonist-specific manner. Crucially, the product A_F_ × P_IA_, which determines the total number of activated receptors, scales directly with cAMP signaling efficacy ratio (correlation coefficient of 0.99) suggesting spatial multimodality underpins ensemble-average efficacy of cellular signaling (New Fig. 5i).). These findings combined reveal the pivotal role multimodality plays in GPCR pharmacology and, ultimately, cellular function.

## Discussion

The plasma membrane is a shared medium, analogous to an optical fiber, through which a plethora of transmembrane protein-based signaling pathways are simultaneously transmitted^49^. This extraordinary capacity to multiplex signals is thought to emerge from the ability of the plasma membrane to organize different protein and lipid species at the nanoscale^25,50,51^. However, it has proven very difficult to directly observe plasma membrane nanodomains and their putative functional properties^52^. By contrast, the direct observation of GPCRs in distinct cellular organelles has been more accessible, and their contribution to so-called ‘location bias’ is thus much better understood^53,54^. Here, we developed a method that leverages 3D live-cell imaging of GPCRs and three different kinds of conformational biosensors^13,14,26^ to map receptor density and intrinsic activation probability across the undulating surface of the plasma membrane (Figs. 2 and 5). Our measurements revealed hitherto elusive nanodomains of strikingly different intrinsic activation probability, a discovery which we refer to as ‘spatial multimodality’.

Spatial multimodality in intrinsic activation probability suggests that, in principle, the same receptor, bound to the same ligand, changes its effective affinity for its signaling partner proteins as it diffuses through different areas of the plasma membrane. This occurs because local nanoscale heterogeneities in the structure and composition of cellular membranes can allosterically bias^34,35^ the conformational equilibria of GPCRs. Spatially heterogeneous membrane properties thus emerge as a fourth operative component of the ternary ligand/GPCR/effector complex^55^. Notably, within such a *quaternary complex* model, intrinsic efficacy is not a constant but a spatiotemporal function ε *(x, y, z, t)*, which assumes a large range of values and thus multiplexes the response of any ternary complex in space and time^56^ (Fig. 6). An intriguing possibility that ought to be examined in the future is that nanodomains may have additional, higher order, functions e.g. driving distinct pathways.

Several key mechanistic insights emerge from this study that upend conventional wisdom. First, the contrasting functional properties of GPCRs in nanodomains do not result from increasing the local density of signaling partners as commonly assumed^25,32,57^, but rather from directly regulating the intrinsic receptor activation probability. Spatially heterogeneous intrinsic activation has been hypothesized^32,52,58^ but, to the best of our knowledge, not directly observed. Second, the mechanism of nanodomain-based regulation of β1AR intrinsic activation does not appear to necessitate changes in the local lipid composition^37^, but rather subtle changes in the local membrane curvature (Fig. 4). Nevertheless, since membrane curvature also appears to influence the spatial organization of negatively charged lipids^38,39^ and is indirectly correlated with cholesterol (Extended Data Fig. 10), it is not unlikely that heterogeneous membrane composition can contribute to the multimodality of other receptors. Third, because plasma membrane curvature can locally persist much longer than lipid or protein diffusion, the lifetime of the nanodomains in unperturbed cells can be 2-5 orders of magnitude larger than proposed by the raft model (∼100s vs ∼ms)^32,52^, which makes it comparable and compatible with the duration of many signaling processes^59^.

For several reconstituted GPCRs, the direct observation of active conformations in the absence of agonists has been a crucial argument in support of the hypothesis that ligands select between preexisting conformations instead of inducing new conformations upon binding (i.e., the ‘conformational selection’ model instead of the ‘induced fit’ model)^4,8^. Our data, however, suggest that only a minor fraction of receptors at the plasma membrane of live cells (1% - 30% depending on the receptor, Fig. 5e) exhibits spontaneous activation. On the one hand, this directly confirms the spontaneous manifestation of active states at the plasma membrane, thus supporting the conformational selection model. On the other hand, it suggests the majority of receptors that do not exhibit detectable spontaneous activation (but activate fully after ligand addition, Fig. 2d) may actually follow the induced fit model. Because the conformational selection model currently underpins our mechanistic understanding of ligand efficacy, further live-cell measurements and quantitative analysis^60^ are warranted to disentangle the putative contribution of the two models/mechanisms to GPCR activation at the plasma membrane.

More broadly, since the spatial multimodality observed here for GPCRs arises from universal physicochemical principles, we expect it will likely influence other transmembrane receptors. Given the cornucopia of membrane curvatures present in all cell types in vivo (plasma membrane, endosomes, endoplasmic reticulum, Golgi apparatus, filopodia, cilia, axons etc.^20,61^), we anticipate that curvature-induced spatial functional multimodality is in general present and underpins signaling in vivo.

## Methods

### Cell lines

Human embryonic kidney (HEK293) cells (ATCC®CRL-1573™) were cultured in DMEM supplemented with 10 % FBS. African green monkey kidney (COS-7) cells were a kind gift from Kenneth Lindegaard Madsen (University of Copenhagen) and were cultured in DMEM supplemented with 10 % FBS. Cell lines were tested routinely for mycoplasma by Eurofins Genomics Mycoplasmacheck. All cell lines were grown at 37 °C, 5 % CO_2_, in an atmosphere with 100 % humidity.

### Cell transfection

All cell lines were grown in 8-well Ibidi® chambers with glass bottoms, where ∼40.000 cells were seeded ∼24 h prior to transient transfection to reach ∼60 % confluency. HEK293 and COS-7 cells were grown on plain glass in an 8-well Ibidi® chamber. For each well, a solution of plasmid, Lipofectamine™ LTX Reagent with PLUS™ was made according to manufacturers’ protocol in the ratio of 1:3:1, and OptiMEM was added to a final volume of 25 μL. The amount of plasmid used for each well was 0.25 μg SNAP-β1AR and 0.186 μg of GFP-miniG_s_ or Nb80-GFP, 0.25 μg SNAP-β2AR and 0.186 μg of GFP-miniG_s_ or Nb80-GFP 0.25 μg, 0.25 μg SNAP-β1AR and 0.186 μg of GFP-actin, 0.25 μg SNAP-GLP1R and 0.186 μg HALO-miniG_s_, 0.25 μg SNAP-GIPR and 0.186 μg HALO-miniG_s_, 0.25 μg SNAP-DRD1 and 0.186 μg GFP-miniG_s_ and 0.25 μg SNAP-dLight1.3b-cpGFP, 0.25 μg SNAP-β1AR and 0.186 μg of GFP-PLCδ (PIP2 biosensor), 0.25 μg SNAP-β1AR and 0.186 μg of GFP-LactC2 (PS biosensor). After transfection, the cells were left to grow for about 16 hours before imaging.

### Live-cell protein labelling

Prior to imaging of GLP1R and GIPR samples, HALO-tagged miniG_s_ was labeled using HALO-JaneliaFluor595 (HALO-JF595) (Promega) by incubating cells in ∼2 μM of HALO-JF595 in DMEM supplemented with 10% FBS for 30 min at 37 °C. Next, cells were washed with Leibovitz’s medium and receptors were labeled with SNAP649 according to manufacturers’ protocol. Briefly, the cell medium was removed from each well and 100 μL of new medium premixed with 0.5 μL of a 50 nmol/μL solution of SNAP-Surface® was added to the cells and the labeling reaction proceeded for 5 min at RT. Similarly, SNAP-tagged β1AR and β2AR were labeled with SNAP649 according to manufacturers’ protocol. Labeling the cells with CellMask was done by adding 20x dilution to the cells for ∼1 min followed by 3x washing with Leibovitz’s.

### Ligands

For β1AR, we added agonist ISO (10 μM), solubilization in Leibovitz’s medium, to HEK293 cells expressing SNAP-labeled β1AR. As ISO is known to hydrolyze, it was stored in powder form under vacuum until usage. For β1AR inverse agonists atenolol (Sigma) and CGP-20712A (Sigma), we added saturating concentrations of 218 nM and 15 nM dissolved in Leibovitz’s medium, respectively. For β1AR, salbutamol (Sigma) and norepinephrine (Sigma) were For DRD1, we added 10 μM of dopamine in Leibovitz’s medium. For the GLP1R we added 500 nM peptide agonist [Aib8]-GLP1(7-36)-Alexa488 to HEK293 cells expressing SNAP-labelled GLP1R. For the GIPR we added 500 nM peptide agonist GIP(1-42) to HEK293 cells expressing SNAP-labelled GLP1R. All peptides were stored in DMSO and diluted in Leibovitz’s medium. All receptor agonists were incubated for 5-10 min before measuring.

Methyl-β-cyclodextrin (MβCD, Sigma) was dissolved in DMEM, and 5 mM was added to HEK293 cells expressing SNAP-β1AR and GFP-miniG_s_ for 10 min at 37 °C. Hereafter, cells were washed with Leibovitz’s medium, labelled with SNAP-tag (as described above) and imaged.

### Live-cell microscopy

Imaging was performed on an Abberior Expert Line system with an Olympus IX83 microscope (Abberior Instruments GmbH) using the Imspector Software v16.3. For imaging SNAP-Surface649, HALO-JF595, cpGFP and GFP, we used respectively 640 nm, 561 nm or 488 nm pulsed excitation laser; fluorescence was detected between 650-720 nm, 580 - 630 nm or 500-550 nm, respectively. For 3D STED imaging we used a pulsed STED line at 775 nm. All XZY stacks were recorded by piezostage (P-736 Pinano, Physik Instrumente, Germany) scanning using a voxel size of 30 nm (dx=dy=dz=30 nm), unless otherwise stated. We used a UPlanSApo x100/1.40 oil immersion objective lens and a pinhole size of 1.0 Airy units (i.e., 100 μm). 3D STED imaging was performed using the easy3D STED module in combination with the adaptive illumination module RESCue^62^. Alignment of the STED and confocal channels was adjusted and verified on Abberior auto-alignment sample, whereas bead measurements were performed with Abberior far-red 30 nm beads. All measurements were made at room temperature and acquired in confocal imaging mode, except when stated otherwise.

### Data acquisition

To map GPCR activation probability, we acquired XZY stacks of the adherent plasma membrane of living cells, as described in detail in ref. ^20^. XZY stacks were on average 6×3×6 μm in size and a single stack was acquired per cell. For experiments that include the addition or titration of ligands, the ‘apo’ and ‘ligand’ conditions were measured on the same cell. Each experimental replicate contains at least 6 different cells and consists of >240.000 data points. While the acquisition of the entire XZY stack takes approximately 1 minute, the temporal resolution of our approach is set by the time required to image a diffraction-limited region in XZ and corresponds to 6 seconds ^20^.

### Reconstructing super-resolved 3D topography and receptor density maps

Membrane topography and receptor density are obtained by image processing of XZY stacks of the adherent plasma membrane using custom written software in MATLAB2019a (The MathWorks®), as described previously^20^. Briefly, the axial (z) point-spread function for every (x,y)-position is fitted with a Gaussian expression. The position of the peak of the Gaussian fit reveals the Z-position of the plasma membrane. The amplitude, i.e. maximum intensity, of the Gaussian fit is proportional to the density of the protein in each (x,y)-pixel. This allows us to, simultaneously, obtain a super-resolved topography map of the adherent plasma membrane of living cell and to measure relative GPCR density using a single fluorescent label, i.e., the fluorescently tagged receptor.

### Extraction of biosensor signal from 3D image stacks

Next to receptor channel, we simultaneously record an XZY stack of the biosensor (miniG or Nb80) channel. We leverage our ability to accurately measure Z-position to extract the relative density of the biosensor at the plasma membrane (Extended Data Fig. 1a). This process is described in detail in the Supplementary Methods. Briefly, we developed a physical model that describes the biosensor intensity along the Z-axis, which consists of a cytosolic contribution and a receptor-bound contribution. These are described by a sigmoidal and Gaussian-shaped curve, respectively (Extended Data Fig. 1b and c). First, all Z profiles were classified for the presence or absence of a Gaussian-like contribution. The intersect-at-half-maximum, or apparent Z-position (Z_app_), was calculated for every trace. A left shift of Z_app_ with respect to the actual Z-position of the plasma membrane, reveals the presence of receptor-bound biosensors (Extended Data Fig. 1c). Using CellMask with miniG as a control, a threshold was determined for the classification of traces (Extended Data Fig. 1e and 1f). Hereafter, all intensity traces with a Gaussian-like contribution were fitted to Equation 1 (Eq. 1):

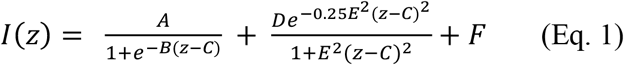

The first part Eq. 1 represents the cytosolic contribution, while the second part is the receptor-bound contribution, which for our optical system is described best by a Lorentzian^63^. A is the amplitude of the sigmoid, B is related to the slope of the sigmoid, C is the position of the membrane, D is the amplitude of the Lorentzian, E is related to the width of the Lorentzian and F is the offset of the curve.

As a result, we obtain the receptor-bound signal (D). Using the error metrics from the fits we filtered out poor fits based on R^2^ (Extended Data Fig. 1g) and uncertainties on the D value. Finally, a 3×3 moving average filter was used to denoise the extracted biosensor intensity maps. Maps of intrinsic activation probability were calculated as the ratio between the extracted biosensor signal and receptor density, as shown in Figs. 2a, 3 and 6. As a result, we obtain a bimodal distribution of activation probabilities at the plasma membrane (Fig. 2d). Notably, the lowest measured P_IA_ in nanodomains (i.e., the left edge of the right hand-side population) is far above the detection limit confirming the histogram is not truncated by low S/N.

To calculate miniG_s_ recruitment in Fig. 1d and e, the average denoised miniG_s_ intensity was calculated for cells where in more than 20% of the pixels miniGs recruitment was detected. This allows us to accurately detect the effects of inverse agonists on basally active cells.

### Measurement of basal and ligand-induced active conformations using dLight1.3b

We measured spatially heterogenous patterns of basal active conformations at the plasma membrane in a biosensor-independent manner by using SNAP-dLight1.3b-cpGFP. An XZ image of the SNAP-channel was recorded in 3D STED mode with a voxel size of 30 nm (dx=dy=dz=30 nm), whereas the cpGFP channel was recorded in confocal mode with dz=150 nm and dx=dy=60 nm. These XZ images were computationally overlaid and the activation probability of dLight1.3b was calculated as the ratio of cpGFP over SNAP.

XZY stacks of SNAP-dLight1.3b-cpGFP were recorded in confocal imaging mode for dLight1.3b experiments with dopamine. The SNAP-channel was recorded with a voxel size of 30 nm (dx=dy=dz=30 nm), whereas the cpGFP channel was recorded with dz=150 nm and dx=dy=60 nm. XZY stacks were denoised using NDSafir^64,65^.

### Measurements of actin density and intrinsic activation probability

We simultaneously acquired *xzy* stacks of cells expressing GFP-actin, HALO-miniG_s_ and SNAP-β1AR. Intrinsic activation probability was measured as described above. Using the Z position of the plasma membrane recovered from the β1AR stack^20^, we calculated actin intensity along the membrane by taking the mean of an 8-pixel average centered along the obtained membrane topography. A threshold was set to the obtained actin density map to identify low and high actin density regions, defined by the lowest and highest 20% of pixels, respectively. For these pixels the average intrinsic activation probability and mean membrane curvature were calculated. Extended Data Fig. 8 reveals that actin density does not affect the intrinsic activation probability, but influences membrane curvature.

### Measurements of PIP2 and PS density

We simultaneously acquired *xzy* stacks of cells expressing SNAP-β1AR and, either GFP-PLCδ (PIP2 biosensor) or GFP-LactC2 (PS biosensor). Similar to measurements of actin density, we used the Z position of the plasma membrane recovered from the β1AR stack^20^ and took an 8-pixel intensity average centered along the obtained membrane topography. This allowed to us obtain PIP2 or PS density at the plasma membrane.

### Fluorescent recovery after photobleaching measurements

To measure the diffusion coefficient of β1AR at the plasma membrane, we used fluorescent recovery after photobleaching (FRAP). A squared XY region of interest (ROI) of 4x4 μm was completely bleached and signal recovery was measured for an XZ slice in the middle of the ROI for 200 s with 5 s intervals. Calculation of the diffusion coefficient for a squared ROI was performed, as previously reported^66^. We calculated the characteristic diffusion time to travel in/out of nanodomains and background using Brownian 2D diffusion at the plasma membrane, where *t* = < *r*, >/4*D*. For a domain of 1 μm in diameter the characteristic diffusion time is ∼1 second. Therefore, for patterns that are stable over ∼7 minutes, receptors will have diffused >400 times over such an area.

### Molecular field theory

The MFT model was developed in Fortran 77 and is described in detail in Supplementary Methods. We used the inactive and active structure of β1AR with accession codes PDB: 2YCW and PDB: 6H7J, respectively.

## Acknowledgements

This work was supported by the Novo Nordisk Foundation (Grant NNF17OC0028176) (D.S.). We also acknowledge funding from the European Union’s Horizon 2020 research and innovation program (Grant agreement No. 891959) (T.P.). We are indebted to B. Kobilka (Stanford University) for providing plasmids, reconstituted β2AR and insightful criticism throughout the project. We are grateful to N. Lambert (Augusta University) for HALO-miniG_s_ plasmid, B. Jones (Imperial College London) for SNAP-GIPR plasmid and K. L. Madsen (University of Copenhagen) for COS-7 cells. We are also thankful to Kirstin Meyer and Orion Weiner (UCSF) for their help with NDSafir.

## Author contributions

D.S. conceived the strategy and was responsible for project management and supervision; G.K. and D.S. designed research; G.K. developed data analysis pipeline with help from L.L. and C.G.S.; G.K. and J.C. performed experiments with help from E.K. and L.L.; M.U. performed all theoretical calculations; A.T. performed β2AR vesicle experiments. S.G.F.R. reconstituted receptors in vesicles; P. K. performed GLP1R and GIPR experiments; A.B. performed β2AR experiments in cells; L.D. and T.P. made constructs and provided help with DRD1 and dLight1.3b experiments; T.W. and C.G.T made constructs and provided help with mutagenesis; G.K. analysed all data; D.S. and G.K. wrote the main text with contributions from C.G.T.; G.K. and D.S. prepared all figures. All authors discussed the results and commented on the manuscript.

## Supplementary information

is available for this paper.

## Data availability

The structure of the inactive and active β1AR are available from the Protein Data bank with accession code 2YCW and 6H7J, respectively. Source data will be provided for this paper.

