## Supplementary material for "Multimodal intrinsic activation of GPCRs in ultrastable plasma membrane nanodomains": Supplmentary Figures and Information

#### Code availability

Source code for extracting density of conformational probe is available on: https://github.com/gkockelkoren/Multimodality. Algorithms for specific custom analysis code are described in detail in the Methods and available upon reasonable request.

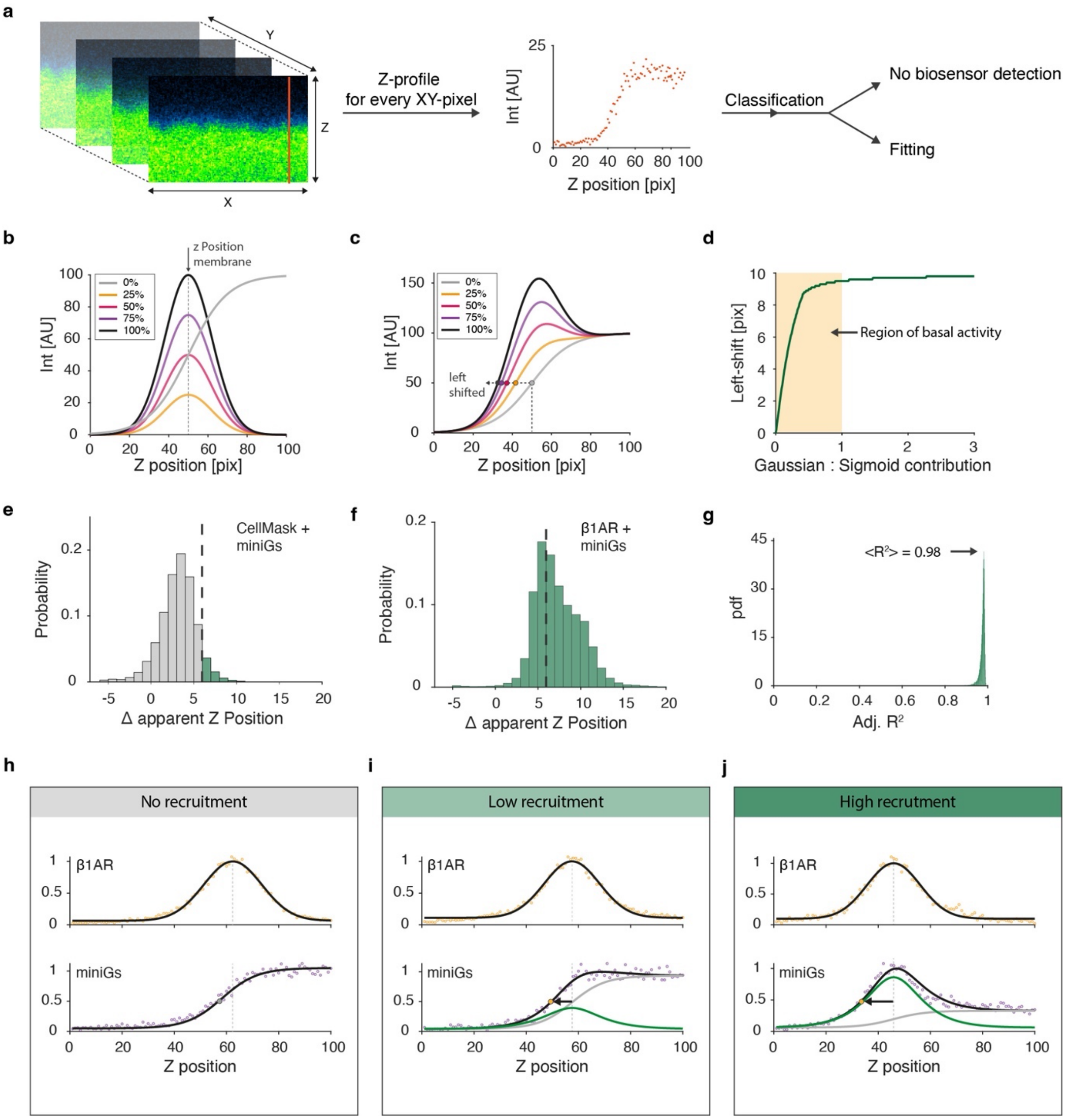


**Extended Data Figure 1. Quantitative extraction of relative miniG density at the plasma membrane of living cells. a,** Schematic pipeline for the extraction of relative miniG density. An XZY stack is recorded by imaging confocal XZ slices of GFP-miniG$_s$ in HEK293 cells and scanning in the Y direction (left). The intensity profile along the Z axis is obtained for every XY-pixel (orange line) (middle). Each intensity profile is classified into 1) undetected or 2) detected miniG at the plasma membrane. For the latter, each profile is fitted with a physical model that extracts miniG intensity. **b,** The XZ intensity profile represents the sum of a sigmoid (cytosolic fraction of miniG) and a Gaussian (receptor-bound miniG). In the absence of receptor-bound miniG at the plasma membrane, the sigmoid is centered at the Z-position of the plasma membrane. **c,** Sum of sigmoid (grey) with increasing amounts of Gaussian contribution (from (b)) is plotted against position on Z-axis. A direct consequence of the Gaussian contribution is a left-ward shift of the apparent Z-position with respect to the Z-position of the membrane. **d,** Magnitude of the left-ward shift (Z-apparent – Z-position) plotted against Gaussian:Sigmoid contribution. At basal state (yellow region, where Gaussian:Sigmoid <1), the left-ward shift is a very sensitive readout of receptor-bound miniG. **e,** Histogram of left-ward shift for negative control CellMask and GFP-miniG$_s$. A threshold for the left-ward shift (dashed line) is determined from the negative control and used to classify traces for the absence or presence of receptor-bound miniG. **f,** Histograms of left-ward shift for SNAP-β1AR and GFP-miniG$_s$. Dashed line indicates the threshold as determined in (e). Cells expressing β1AR exhibit a larger left-ward shift than the CellMask-control. **g,** After classification, all traces with a Gaussian component are fitted with the sum of a Gaussian and sigmoid. Histogram of goodness of fit, calculated as adjusted $R^2$, reveals high-quality fits close to 1. **h, i, j,** Representative examples of XZ traces for the scenarios of 'no recruitment', 'medium recruitment' and 'high recruitment' of miniG from left to right. Top profiles represent XZ traces of the β1AR from which we extract receptor density and membrane position in Z (shown as dashed grey line), as described in ref. [20]. Bottom profiles are XZ intensity profiles from the miniG channel that are fitted with either with a sigmoid (grey), or sigmoid (grey) and Gaussian (green). The black arrow indicates the left-ward shift of the trace due to the presence of a Gaussian. All data is representative of CellMask, $n_C$ = 18 and $n_R$ = 3, and of β1AR, $n_C$ = 82 and $n_R$ = 11.

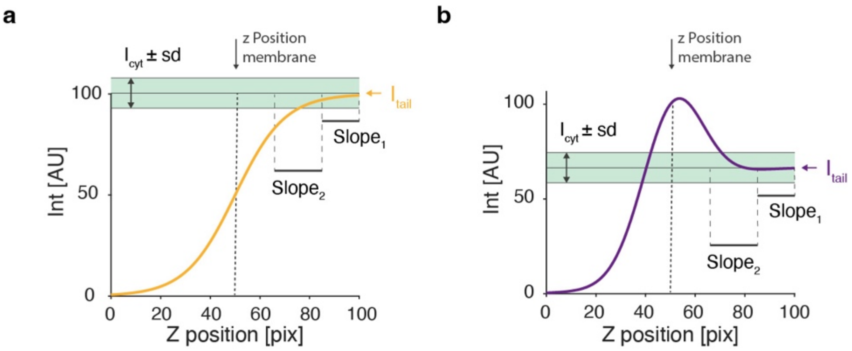


**Extended Data Figure 2. Additional parameters considered for classification criteria of intensity traces. a, b,** The presence of receptor-bound miniG at the plasma membrane is primarily detected through a left-ward shift of the XZ intensity profile with respect to the Z-position of the membrane (Extended Data Fig. 1). We have added additional classification criteria to correct classification of traces. Schematic examples of intensity traces along the Z-axis of no recruitment (a) and recruitment (b). Next to the left-ward shift of the trace with respect to the Z position of the membrane, 3 other parameters are considered to classify traces: Slope1, Slope2 and Icyt (see detailed description in Supplementary Information).

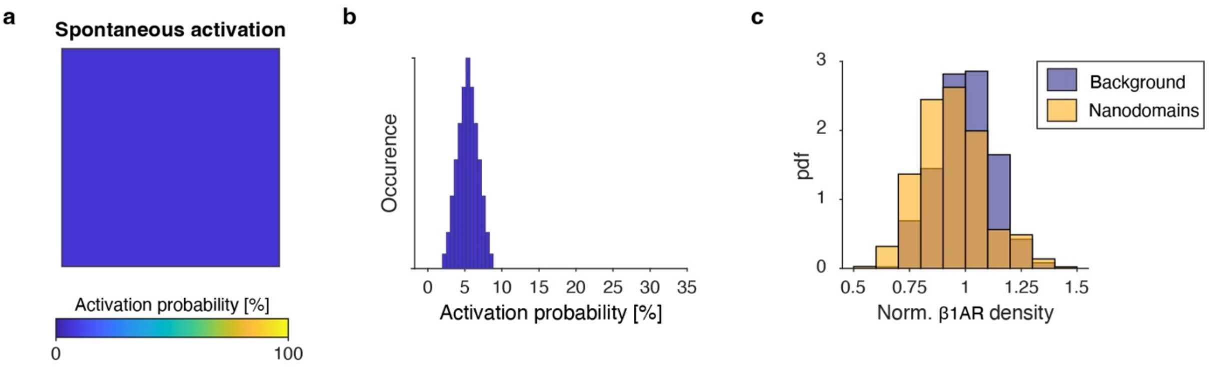


**Extended Data Figure 3. Basal activation probability of receptors is not uniform at the plasma membrane. a,** Traditionally, GPCR basal signaling is thought to rely on spontaneous (i.e., thermal energy induced) sampling of active conformations. Consequently, it is believed that all receptors are equally probably to sample an active conformation at the plasma membrane. Accordingly, we would expect to observe a uniform spatial map of activation probability. As shown in Fig. 2, this is not the case and a map of intrinsic activation probability reveals evident spatial heterogeneity. **b,** The distribution of activation probabilities for the map shown in (a) is represented by a narrow normally distributed histogram. **c,** Histograms of β1AR density (for the area as shown in Figs. 2a and b) show that receptors are present in both nanodomains and background. Thus, 0% activation probability in the background is not due to the absence of receptors in these areas.

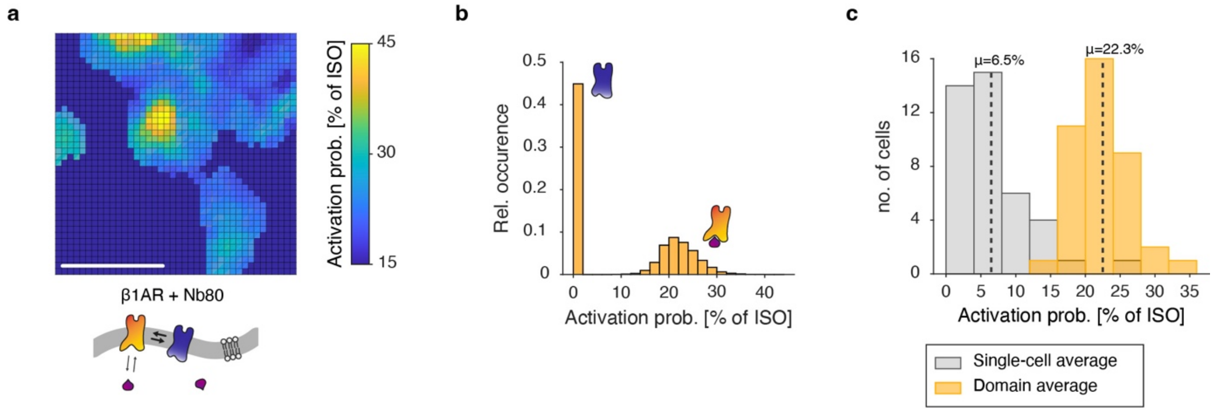


**Extended Data Figure 4. Nanobody-80 reveals quantitively comparable spatial patterns of basal activation probability to miniG$_s$. a,** Map of basal activation probability for β1AR as measured by nanobody-80 (Nb80). Color-scale is normalized to activation by ISO (i.e., 100%). Cartoon below the map represents the experimental setup. Scale bar, 500 nm. **b,** Histogram of spatially resolved activation probability normalized to ISO. Like miniG$_s$, the activation probability shows a bimodal distribution. **c,** Histograms of average activation probability (as a % of full activation by ISO) for the entire cell (i.e., domains and background) (grey) and for domains only (yellow). Dashed lines indicate means of each population. Single-cell and domain average for Nb80 are very similar to miniG$_s$ (c.f. 3.9% and 23.9%, Extended Data Fig. 3). Data is $n_C = 41$ and $n_R = 7$.

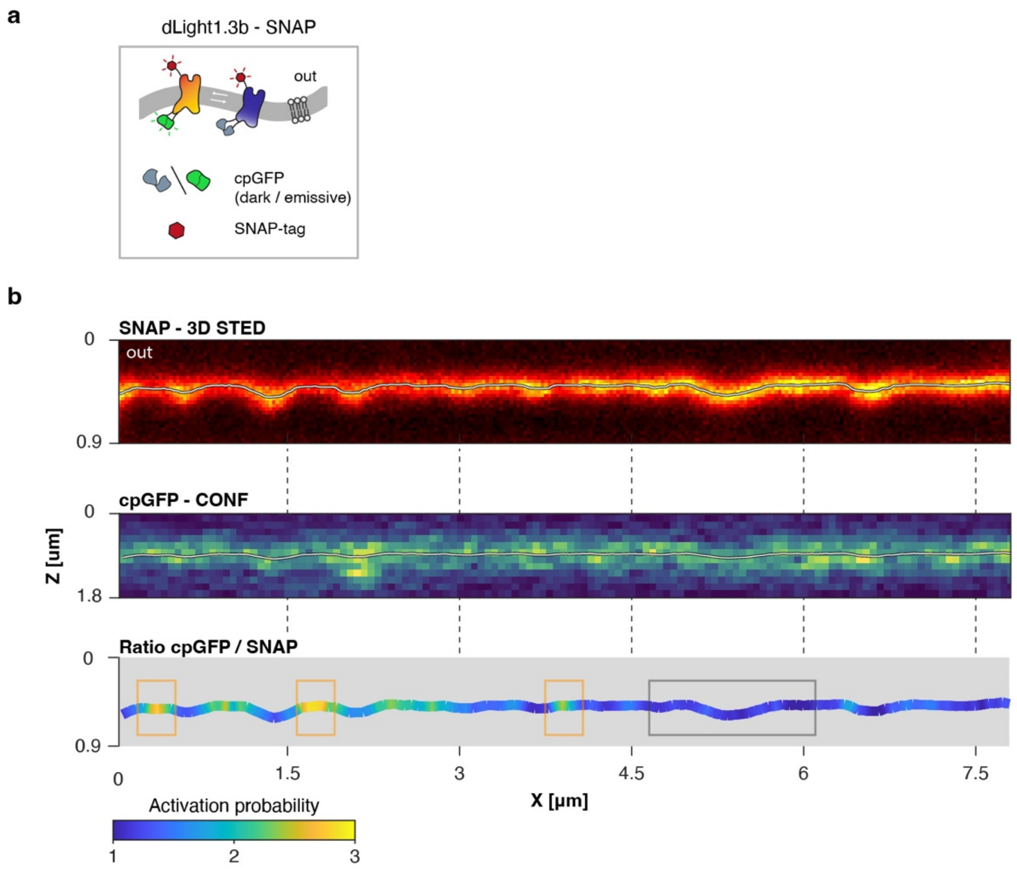


**Extended Data Figure 5. Ratio-metric imaging of SNAP-dLight1.3b reveals spatially heterogeneous distribution of intrinsic activation probability. a,** Cartoon illustrating how activation of the DRD1 analog, SNAP- dLight1.3b, results in fluorescence emission by cpGFP. **b,** XZ micrographs of SNAP- dLight1.3b are shown. Top: SNAP channel imaged in 3D STED imaging modality. Middle: cpGFP channel imaged in confocal mode. Bottom: Ratio of cpGFP / SNAP using the middle and top channel. Yellow boxes outline basal active nanodomains and black box outlines a region of low/zero activation probability. Data is representative of $n_C = 34$ and $n_R = 3$.

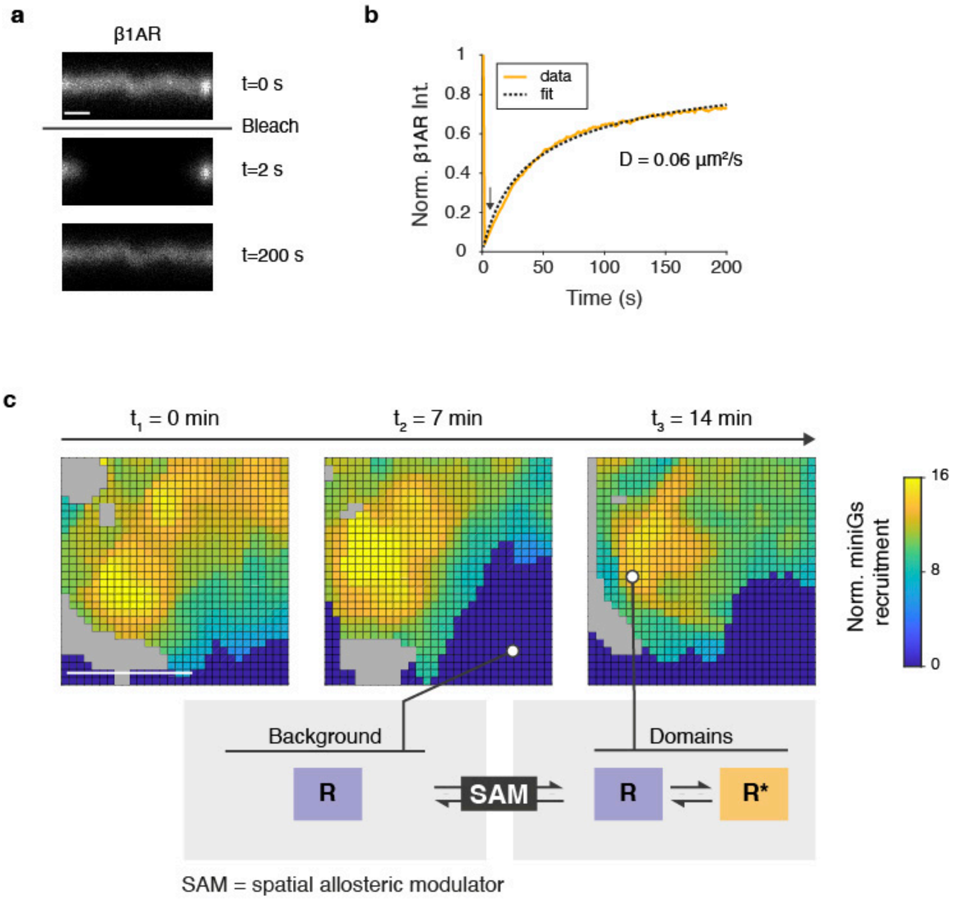


**Extended Data Figure 6. Measurements of characteristic timescales of β1AR diffusion and nanodomain stability at the plasma membrane. a,** Micrographs show fluorescent recovery after photobleaching (FRAP) of confocal XZ-slices of the adherent part of the plasma membrane expressing β1AR. Micrographs are acquired prior to bleaching (top), immediately after bleaching (middle) and after 200 s (bottom). All images have identical color scales. Data is from $n_R$ = 3. Scale bar, 1 μm. **b,** FRAP curve (in yellow) shows β1AR intensity (normalized to the intensity before photobleaching) versus time. Black arrow indicates moment of bleaching. Dashed line represents the fit to the data and allows calculation of the reported diffusion coefficient (D=0.06 $\mu m^2/s$). **c,** Maps of miniG$_s$ recruitment at 3 time points separated by 7-minute intervals reveal stability of nanodomains over time. Grey areas represent regions that have been filtered out (see Methods). Scalebar for all maps, 500 nm. Illustration highlights that in dark blue regions the inactive state (R) prevails, while in nanodomains both the inactive and active (R*) state exist. As shown in Fig. 3, the plasma membrane acts a spatial allosteric modulator (SAM).

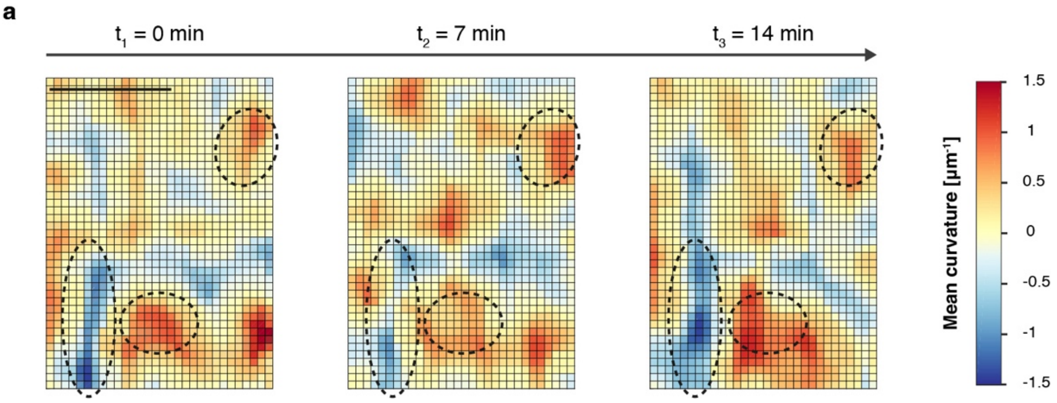


**Extended Data Figure 7. Ultra-long lived membrane curvature features of the plasma membrane persist for periods of up to ~10 minutes. a,** Maps of membrane curvature measured at the same position at 3 time points with 7 minutes intervals. Over this period some features remain stable while others are dynamic. Dashed outlines indicate representative regions of stable membrane curvature. Similar to the basal activity patterns, the curvature of certain topographical features is stable, whereas others are more dynamic. Data is from $n_R$ = 3. Scale bar, 500 nm.

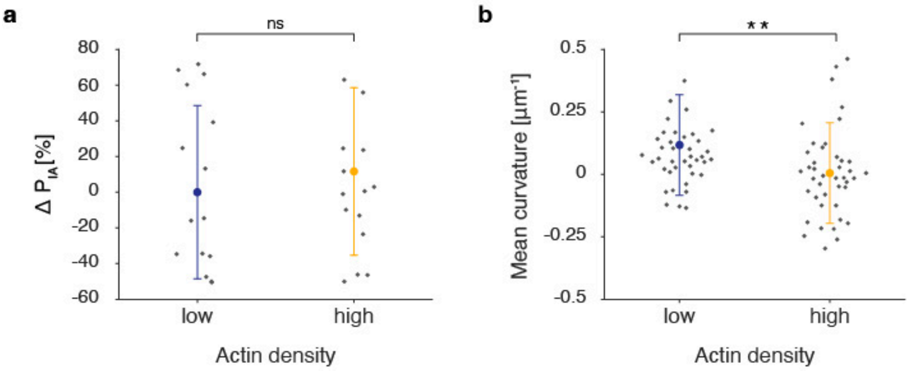


**Extended Data Figure 8. Actin density has no significant effect on $P_{IA}$, but influences membrane curvature. a,** Average change in $P_{IA}$ is calculated for regions with low and high actin density. No significant correlation is observed between $P_{IA}$ and actin density. Data is mean of all cells +/- SD. Data represents $n_C$ = 16 and $n_R$ =2. ns (*P*=0.4941). **b,** Average mean membrane curvature is calculated for regions of low and high actin density. An increase in negative membrane curvature is observed for high actin density. Data is mean of all cells +/- SD. ***P*=0.0076. Data represents $n_C$ = 42 and $n_R$ =6.

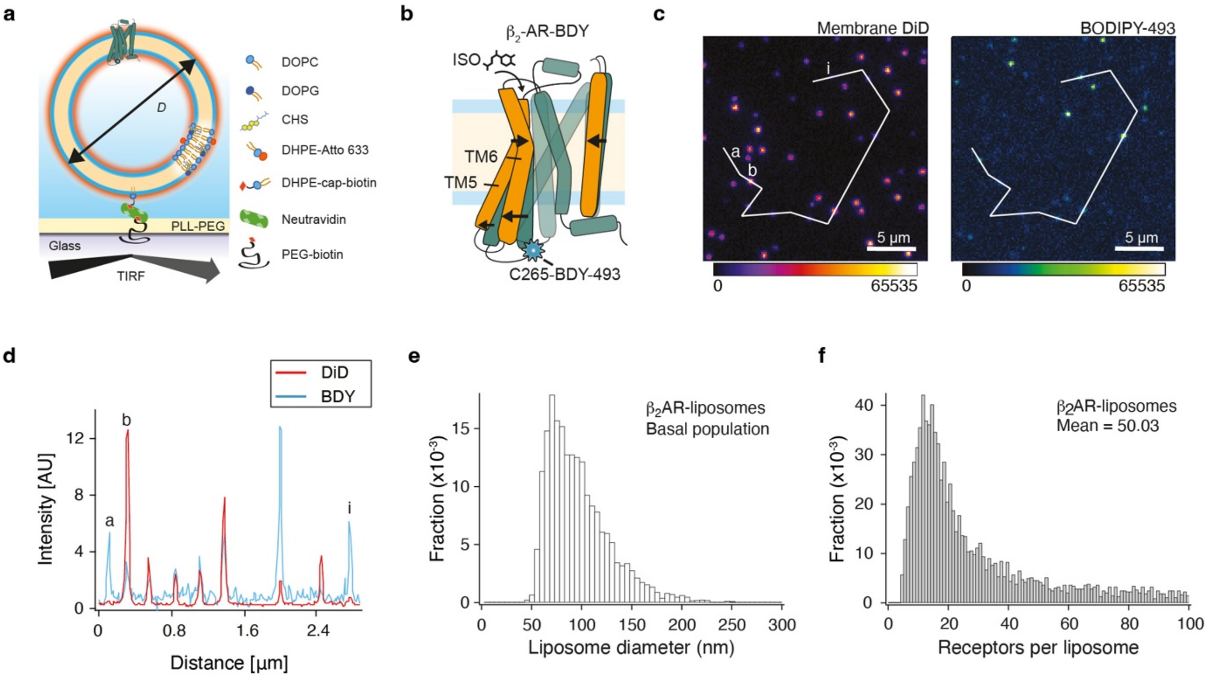


**Extended Data Figure 10. Quantitative fluorescence recording of β2AR conformational changes in liposomes. a,** Immobilized liposomes, reconstituted with $\beta_2$AR-BDY onto a polymer passivated surface via neutravidin-biotin binding, allow quantification of changes in receptor conformation and liposome diameter. **b,** Schematics of the β2AR-BDY labeled with BDY-493 to TM6 at the cysteine C265 (active receptor in orange and inactive in green). Black arrows indicate changes in the conformational state upon receptor activation with a full agonist. **c,** Fluorescent micrographs of an array of liposomes on glass slide show membrane channel (DiD) and BDY channel. **d,** Line profile through liposomes of images in (c) shows that the intensities do not correlate in amplitudes, as smaller liposomes (decreased DiD signal) show increased BDY signal. This is highlighted for the liposomes marked a, b and i in (c). **e,** Distribution of liposome diameters. **f,** Distribution of number of receptors per liposome. Data is representative of $n_R$ = 8.

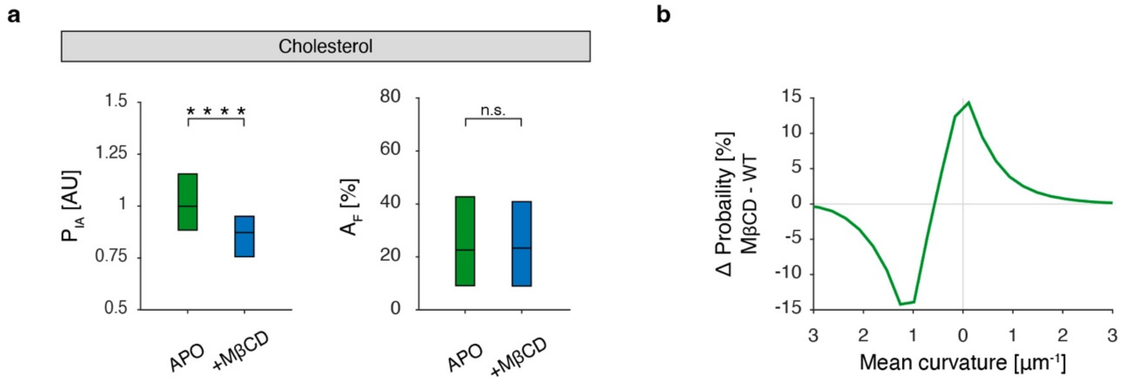


**Extended Data Figure 10. Depletion of cholesterol with methyl-β-cyclodextrin (MβCD) affects $P_{IA}$, likely through an indirect effect on cellular morphology. a,** Boxplots for $P_{IA}$ (c) and $A_F$ (d) show differences between basal β1AR state in HEK293 treated without and with MβCD. Cholesterol depletion decreases $P_{IA}$, but does not affect $A_F$. For $P_{IA}$, ****$P$=4.2×10$^{-4}$, and for $A_F$, n.s. $P$=0.67, with two-sided Kolmogorov-Smirnov test. **b,** Change in probability of observing a certain mean curvature after treatment of cells with MβCD. Cholesterol depletion causes a decrease in probability at negative curvature and an increase in zero curvature areas. For β1AR APO in HEK293 cells, $n_C$ = 82 and $n_R$ = 11, for β1AR APO with MβCD, $n_C$ = 37 and $n_R$ =3.

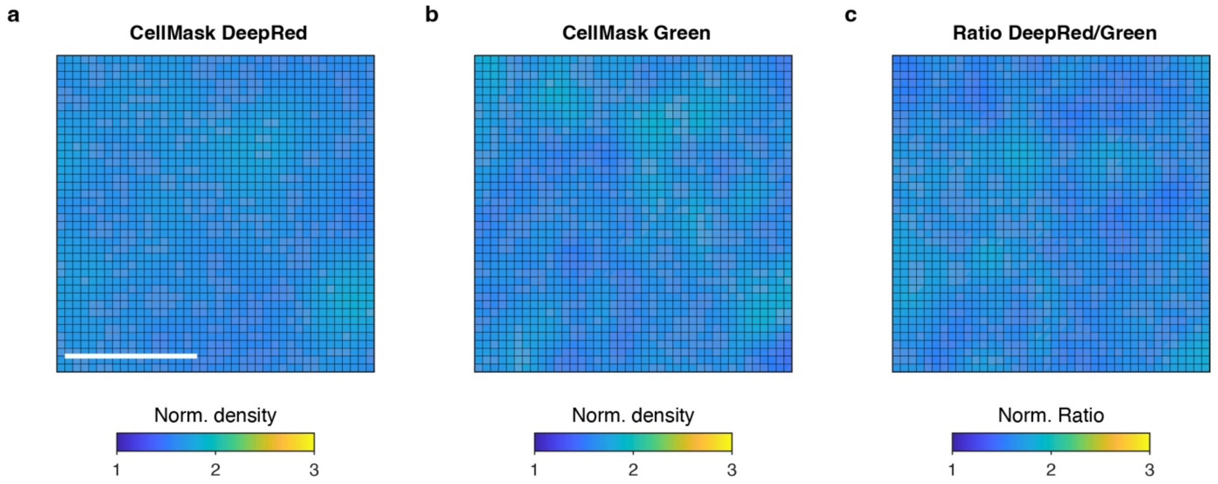


**Extended Data Fig. 11. Ratio-metric analysis of two-color CellMask staining of the plasma membrane reveals a uniform surface. a, b, c,** XY surface density of CellMask DeepRed (a) and CellMask Green (b) and their ratio (c) at the plasma membrane are uniform. This demonstrates the absence of putative imaging and analysis artifacts. Scale bar, 500 nm. Data is representative of $n_C = 8$ cells and $n_R = 2$ experiments.

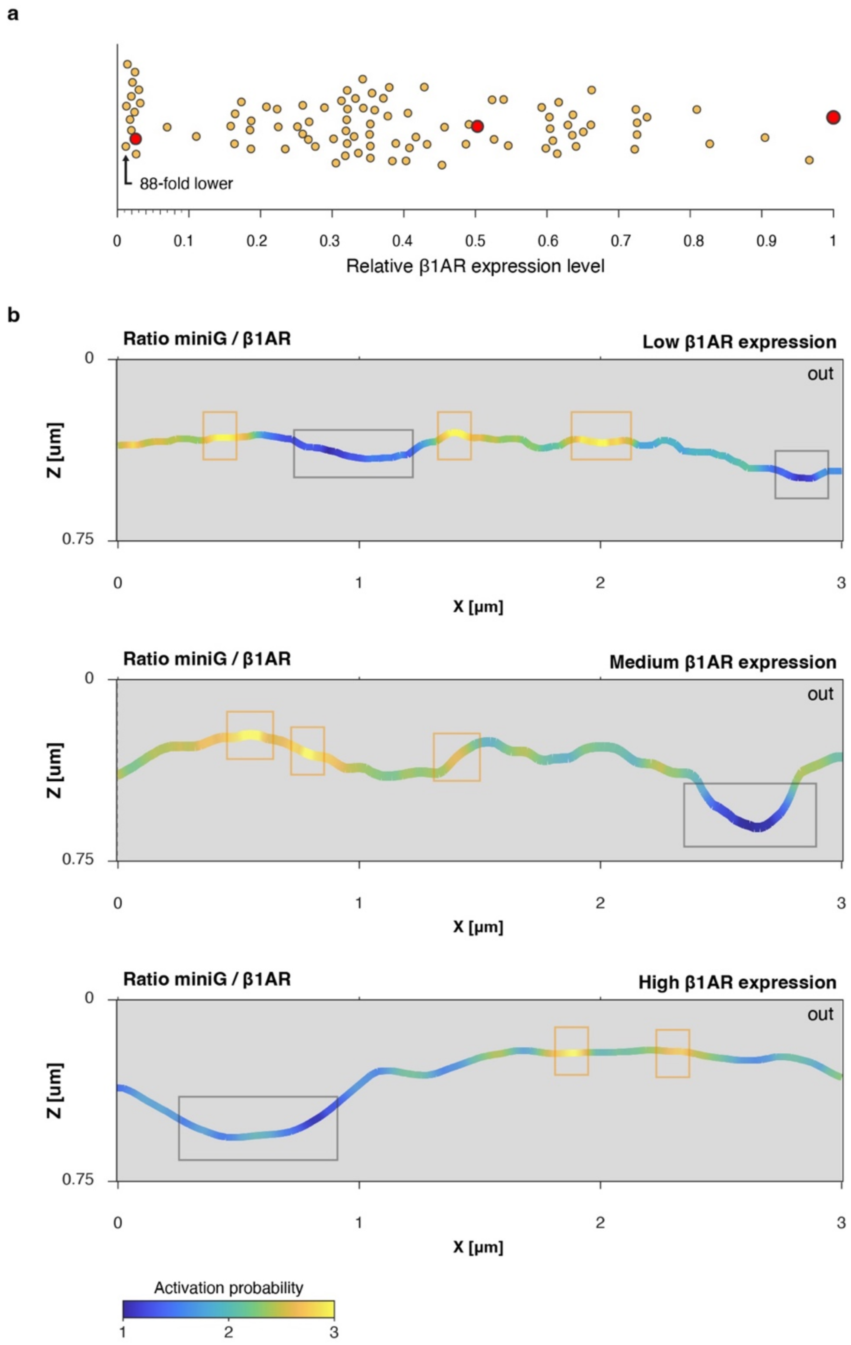


**Extended Data Figure 12. Nanodomains persist at 88-fold lower receptor expression levels. a,** Scatter plot of β1AR expression level measured in experiments. Each data point represents the average expression of one cell. Receptor expression levels span almost 2 orders of magnitude and consistently reveal the existence of nanodomains of contrasting activation probability. **b,** XZ view of the plasma membrane color-coded by the ratio of miniG$_s$/β1AR for a cell expressing 40-fold less β1AR than previously shown. Yellow boxes outline active nanodomains and the black box outlines a region of low activation probability. Data is recorded after activation with a saturating concentration of agonist ISO (8.7 µM). Data is representative of 29 XZ slices from $n_C$ = 19 cells and $n_R$ = 3 experiments.

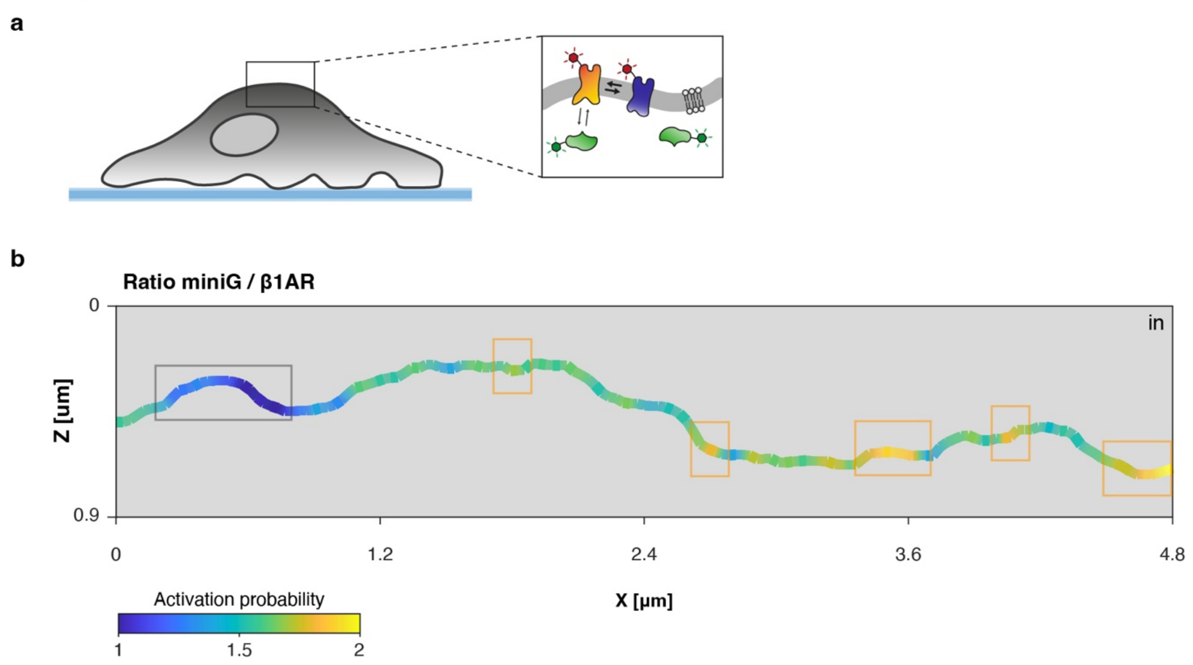


**Extended Data Figure 13. Nanodomains exist on freestanding membranes. a,** Schematic illustrating the existence of segregated active and inactive receptors, i.e., nanodomains of contrasting activation probability, at the freestanding membrane on the top of the cell. **b,** XZ view of the plasma membrane color-coded by the ratio of miniG$_s$/β1AR at the freestanding plasma membrane on the top of the cell. Yellow boxes outline active nanodomains and the black box outlines a region of low activation probability. Data is recorded after activation with a saturating concentration of agonist ISO (8.7 µM). Data is representative of $n_C = 36$ and $n_R = 3$.

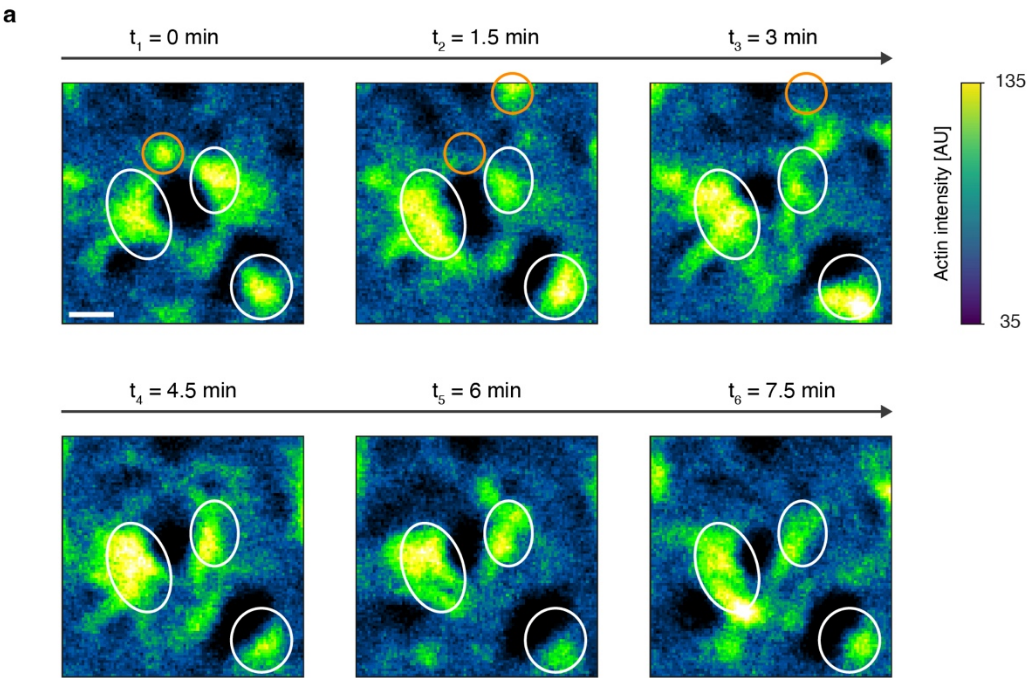


**Extended Data Figure 14. Some actin structures are stable and some are dynamic over 5 minutes. a,** XY images of actin-GFP over time reveal stable (white circles) and dynamic (orange circle) features over timescales longer than 5 minutes. This agrees with our observation that nanodomains of contrasting activation probability are ultra-stable, i.e. persisting for > 5 minutes, yet show dynamics over longer timescales. Data is representative of $n_C$ = 10 cells and $n_R$ = 3 experiments. Scalebar applies to all images, 500 nm.

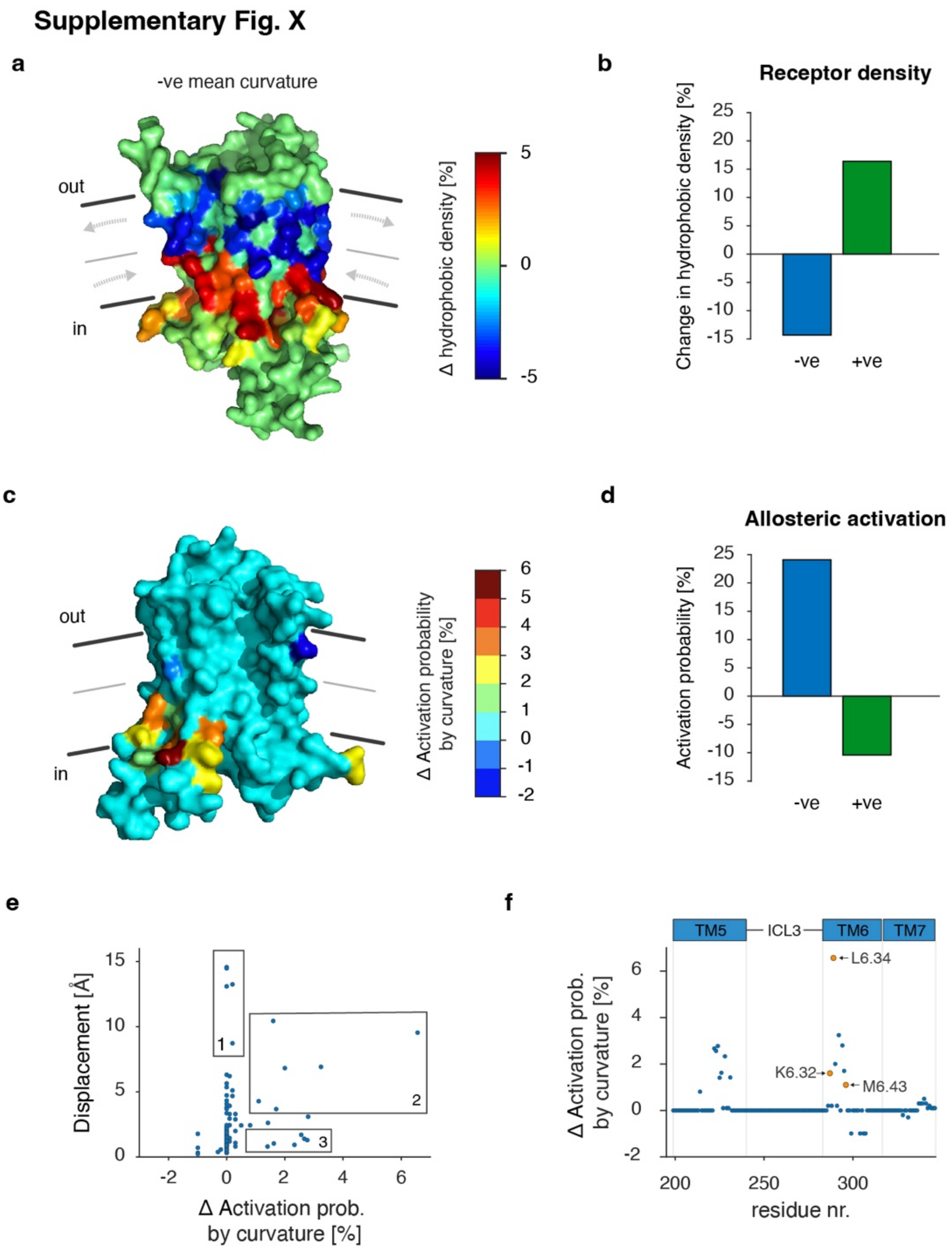


**Extended Data Figure 15. Molecular mapping of membrane curvature-effects on GPCR density and allosteric activation. a,** Color maps of the hydrophobic contribution to receptor sorting (normalized to 0 mean curvature) mapped onto the volume view of β1AR (PDB: 2YCW) at single-residue resolution are shown in the middle and on the right. The receptor is embedded in a membrane with a negative mean curvature of $-1.33\ \mu m^{-1}$. Red residues in the inner leaflet indicate an increase in the hydrophobic contribution, whereas blue residues in the outer leaflet show a decrease in the hydrophobic contribution to the overall curvature coupling. Gray arrows in the bilayer represent the compression/expansion of the intra- and extracellular leaflet as a direct consequence of membrane bending. **b,** Bar graph shows the sum of hydrophobic densities over all amino acids. The sum reflects the increase in receptor density observed at positive curvature and the decrease in density at negative curvature, as shown in ref. [20]. **c,** Color map of the contribution of curvature to receptor activation mapped onto the cartoon view of active β1AR (PDB: 6H7J) at single residue resolution. **d,** Bar graph shows the sum of activation probabilities over all amino acids. The sum reflects the probability of observing the active state at negative or positive curvature. **e,** Scatter plot of amino acid displacement upon activation versus the change in activation probability by curvature. Boxed areas show three scenarios: 1) a small change in activation probability but large displacement, 2) a large change in activation probability and large displacement, and 3) a large change in activation probability but small displacement. **f,** Scatter plot of the change in activation probability for every β1AR residue due to curvature (negative versus positive) in TM5, 6 and 7. Highlighted are three amino acids (K6.32,

L6.34 and M6.43) that have been previously identified to increase receptor activation through classical signaling assays.

#### References


1 Pierce, K. L., Premont, R. T. & Lefkowitz, R. J. Seven-transmembrane receptors. *Nat. Rev. Mol. Cell Biol.* **3**, 639-650 (2002).

2 Wootten, D., Christopoulos, A., Marti-Solano, M., Babu, M. M. & Sexton, P. M. Mechanisms of signalling and biased agonism in G protein-coupled receptors. *Nat. Rev. Mol. Cell Biol.* **19**, 638-653 (2018).

3 Shimada, I., Ueda, T., Kofuku, Y., Eddy, M. T. & Wuthrich, K. GPCR drug discovery: integrating solution NMR data with crystal and cryo-EM structures. *Nat. Rev. Drug Discov.* **18**, 59-82 (2019).

4 Weis, W. I. & Kobilka, B. K. The molecular bases of G protein-coupled receptor activation. *Annual Review of Biochemistry* **87**, 879-919 (2018).

5 Nygaard, R. *et al.* The dynamic process of beta(2)-adrenergic receptor activation. *Cell* **152**, 532-542 (2013).

6 Vafabakhsh, R., Levitz, J. & Isacoff, E. Y. Conformational dynamics of a class C G-protein-coupled receptor. *Nature* **524**, 497-501 (2015).

7 Manglik, A. *et al.* Structural Insights into the Dynamic Process of beta2-Adrenergic Receptor Signaling. *Cell* **161**, 1101-1111 (2015).

8 Ye, L., Van Eps, N., Zimmer, M., Ernst, O. P. & Prosser, R. S. Activation of the A2A adenosine G-protein-coupled receptor by conformational selection. *Nature* **533**, 265-268 (2016).

9 Gregorio, G. G. *et al.* Single-molecule analysis of ligand efficacy in beta(2)AR-G-protein activation. *Nature* **547**, 68-73 (2017).

10 Wingler, L. M. *et al.* Angiotensin Analogs with Divergent Bias Stabilize Distinct Receptor Conformations. *Cell* **176**, 468-478 e411 (2019).

11 Hilger, D. *et al.* Structural insights into differences in G protein activation by family A and family B GPCRs. *Science* **369** (2020).

12 Lerch, M. T. *et al.* Viewing rare conformations of the beta(2) adrenergic receptor with pressure-resolved DEER spectroscopy. *Proc. Natl. Acad. Sci. USA* **117**, 31824-31831 (2020).

13 Irannejad, R. *et al.* Conformational biosensors reveal GPCR signalling from endosomes. *Nature* **495**, 534-538 (2013).

14 Wan, Q. *et al.* Mini G protein probes for active G protein-coupled receptors (GPCRs) in live cells. *J. Biol. Chem.* **293**, 7466-7473 (2018).

15 Warne, T. *et al.* Structure of a beta1-adrenergic G-protein-coupled receptor. *Nature* **454**, 486-491 (2008).

16 Carpenter, B. & Tate, C. G. Engineering a minimal G protein to facilitate crystallisation of G protein-coupled receptors in their active conformation. *Protein Eng. Des. Sel.* **29**, 583-594 (2016).

17 Stoeber, M. *et al.* A genetically encoded biosensor reveals location bias of opioid drug action. *Neuron* **98**, 963-976 e965 (2018).

18 Crilly, S. E., Ko, W., Weinberg, Z. Y. & Puthenveedu, M. A. Conformational specificity of opioid receptors is determined by subcellular location irrespective of agonist. *Elife* **10** (2021).

19 Radoux-Mergault, A., Oberhauser, L., Aureli, S., Gervasio, F. L. & Stoeber, M. Subcellular location defines GPCR signal transduction. *Science Signaling* **9** (2023).

20 Kockelkoren, G. *et al.* Molecular mechanism of GPCR spatial organization at the plasma membrane. *Nat. Chem. Biol.* (2023).

21 Schihada, H., Shekhani, R. & Schulte, G. Quantitative assessment of constitutive G protein–coupled receptor activity with BRET-based G protein biosensors. *Science Signaling* **14** (2021).
22 Ballesteros, J. A. *et al.* Activation of the beta 2-adrenergic receptor involves disruption of an ionic lock between the cytoplasmic ends of transmembrane segments 3 and 6. *J. Biol. Chem.* **276**, 29171-29177 (2001).
23 Baker, J. G., Hall, I. P. & Hill, S. J. Agonist and Inverse Agonist Actions of Human beta2-Adrenoreceptor Provide Evidence for Agonist-Directed Signaling. *Molecular Pharmacology* **64** (2003).
24 Michel, M. C., Michel-Reher, M. B. & Hein, P. A Systematic Review of Inverse Agonism at Adrenoceptor Subtypes. *Cells* **9** (2020).
25 Sungkaworn, T. *et al.* Single-molecule imaging reveals receptor-G protein interactions at cell surface hot spots. *Nature* **550**, 543-547 (2017).
26 Patriarchi, T. *et al.* Ultrafast neuronal imaging of dopamine dynamics with designed genetically encoded sensors. *Science* **360** (2018).
27 Manchanda, Y. *et al.* Expression of mini-G proteins specifically halt cognate GPCR trafficking and intracellular signaling. *BioRXiv* (2021).
28 Ahmed, M. *et al.* Beta-blockers show inverse agonism to a novel constitutively active mutant of beta1-adrenoceptor. *J. Pharmacol. Sci.* **102**, 167-172 (2006).
29 Engelhardt, S., Grimmer, Y., Fan, G. & Lohse, M. J. Constitutive Activity of the Human beta1-Adrenergic Receptor in beta1-Receptor Transgenic Mice. *Molecular Pharmacology* **60** (2001).
30 Milligan, G. Constitutive Activity and Inverse Agonists of G Protein-Coupled Receptors: a Current Perspective. *Molecular Pharmacology* **64** (2003).
31 Yanagawa, M. H., M.; Togashi, Y.; Abe, M.; Yamashita, T.; Shichida, M.; Murata, M.; Ueda, M.; Sako, Y.;. Single-molecule diffusion-based estimation of ligand effects on G protein–coupled receptors. *Science Signaling* (2018).
32 Kusumi, A. *et al.* Dynamic organizing principles of the plasma membrane that regulate signal transduction: commemorating the fortieth anniversary of Singer and Nicolson's fluid-mosaic model. *Annu. Rev. Cell Dev. Biol.* **28**, 215-250 (2012).
33 Maslov, I. *et al.* Sub-millisecond conformational dynamics of the A(2A) adenosine receptor revealed by single-molecule FRET. *Commun Biol* **6**, 362 (2023).
34 Soubias, O., Teague, W. E., Jr., Hines, K. G. & Gawrisch, K. Rhodopsin/lipid hydrophobic matching-rhodopsin oligomerization and function. *Biophys. J.* **108**, 1125-1132 (2015).
35 Brown, M. F. Soft matter in lipid-protein interactions. *Annu. Rev. Biophys.* (2017).
36 Doherty, G. J. & McMahon, H. T. Mediation, modulation, and consequences of membrane-cytoskeleton interactions. *Annu. Rev. Biophys.* **37**, 65-95 (2008).
37 Dawaliby, R. *et al.* Allosteric regulation of G protein-coupled receptor activity by phospholipids. *Nat. Chem. Biol.* **12** (2016).
38 Sejdiu, B. I. & Tieleman, D. P. Lipid-Protein Interactions Are a Unique Property and Defining Feature of G Protein-Coupled Receptors. *Biophys. J.* **118**, 1887-1900 (2020).
39 Thakur, N. *et al.* Anionic phospholipids control mechanisms of GPCR-G protein recognition. *Nat. Commun.* **14**, 794 (2023).
40 May, S. & Ben-Shaul, A. A molecular model for lipid-mediated interaction between proteins in membranes. *Phys. Chem. Chem. Phys.* **2**, 4494-4502 (2000).
41 Sperotto, M. M., May, S. & Baumgaertner, A. Modelling of proteins in membranes. *Chem. Phys. Lipids* **141**, 2-29 (2006).
42 Larsen, J. B. *et al.* Membrane curvature enables N-Ras lipid anchor sorting to liquid-ordered membrane phases. *Nat. Chem. Biol.* **11**, 192-194 (2015).

43 Larsen, J. B. *et al.* Membrane curvature and lipid composition synergize to regulate N-Ras anchor recruitment. *Biophys. J.* **113**, 1269-1279 (2017).
44 Larsen, J. B. *et al.* How membrane geometry regulates protein sorting independently of mean curvature. *ACS Cent. Sci.* **6**, 1159-1168 (2020).
45 Fung, J. J. *et al.* Ligand-regulated oligomerization of beta(2)-adrenoceptors in a model lipid bilayer. *EMBO J.* **28**, 3315-3328 (2009).
46 Mathiasen, S. *et al.* Nanoscale high-content analysis using compositional heterogeneities of single proteoliposomes. *Nat. Methods* **11**, 931-934 (2014).
47 Baker, J. G. The selectivity of beta-adrenoceptor agonists at human beta1-, beta2- and beta3-adrenoceptors. *Br J Pharmacol* **160**, 1048-1061 (2010).
48 Zhang, Y. *et al.* Cryo-EM structure of the activated GLP-1 receptor in complex with a G protein. *Nature* **546**, 248-253 (2017).
49 Grecco, H. E., Schmick, M. & Bastiaens, P. I. Signaling from the living plasma membrane. *Cell* **144**, 897-909 (2011).
50 Pontier, S. M. *et al.* Cholesterol-dependent separation of the beta2-adrenergic receptor from its partners determines signaling efficacy: insight into nanoscale organization of signal transduction. *J. Biol. Chem.* **283**, 24659-24672 (2008).
51 Lohse, M. J. & Hofmann, K. P. Spatial and temporal aspects of signaling by G-protein-coupled receptors. *Mol. Pharmacol.* **88**, 572-578 (2015).
52 Levental, I. & Lyman, E. Regulation of membrane protein structure and function by their lipid nano-environment. *Nat. Rev. Mol. Cell Biol.* **24**, 107-122 (2023).
53 Irannejad, R. *et al.* Functional selectivity of GPCR-directed drug action through location bias. *Nat. Chem. Biol.* **13**, 799-806 (2017).
54 Tsvetanova, N. G. & von Zastrow, M. Spatial encoding of cyclic AMP signaling specificity by GPCR endocytosis. *Nat. Chem. Biol.* **10**, 1061-1065 (2014).
55 De Lean, A., Stadel, J. M. & Lefkowitz, R. J. A ternary complex model explains the agonist-specific binding properties of the adenylate cyclase-coupled beta-adrenergic receptor. *Journal of Biological Chemistry* **255**, 7108-7117 (1980).
56 Grundmann, M. & Kostenis, E. Temporal Bias: Time-Encoded Dynamic GPCR Signaling. *Trends Pharmacol. Sci.* **38**, 1110-1124 (2017).
57 Sezgin, E., Levental, I., Mayor, S. & Eggeling, C. The mystery of membrane organization: composition, regulation and roles of lipid rafts. *Nat. Rev. Mol. Cell Biol.* **18**, 361-374 (2017).
58 Kuriyan, J. & Eisenberg, D. The origin of protein interactions and allostery in colocalization. *Nature* **450**, 983-990 (2007).
59 Huang, W. Y. C. *et al.* A molecular assembly phase transition and kinetic proof reading modulate Ras activation by SOS. *Science* (2019).
60 Hammes, G. G., Chang, Y. C. & Oas, T. G. Conformational selection or induced fit: a flux description of reaction mechanism. *Proc. Natl. Acad. Sci. USA* **106**, 13737-13741 (2009).
61 Zimmerberg, J. & Kozlov, M. M. How proteins produce cellular membrane curvature. *Nat. Rev. Mol. Cell Biol.* **7**, 9-19 (2006).
62 Staudt, T. *et al.* Far-field optical nanoscopy with reduced number of state transition cycles. *Optics Express* **19** (2011).
63 Vicidomini, G. *et al.* STED nanoscopy with time-gated detection: theoretical and experimental aspects. *PLoS One* **8**, e54421 (2013).
64 Kervrann, C. & Boulanger, J. Optimal spatial adaptation for patch-based image denoising. *IEEE Trans. Image Process.* **15**, 2866-2878 (2006).
65 Carlton, P. M. *et al.* Fast live simultaneous multiwavelength four-dimensional optical microscopy. *Proc. Natl. Acad. Sci. USA* **107**, 16016-16022 (2010).

66 Blumenthal, D., Goldstien, L., Edidin, M. & Gheber, L. A. Universal Approach to FRAP Analysis of Arbitrary Bleaching Patterns. *Sci Rep* **5**, 11655 (2015).
67 Samama, P., Cotecchia, S., Costa, T. & Lefkowitz, R. J. A mutation-induced activated state of the beta 2-adrenergic receptor. Extending the ternary complex model. *Journal of Biological Chemistry* **268**, 4625-4636 (1993).
68 Fung, J. J. *et al.* Ligand-regulated oligomerization of beta(2)-adrenoceptors in a model lipid bilayer. *EMBO J* **28**, 3315-3328 (2009).
69 Christensen, S. M., Bolinger, P. Y., Hatzakis, N. S., Mortensen, M. W. & Stamou, D. Mixing subattolitre volumes in a quantitative and highly parallel manner with soft matter nanofluidics. *Nature Nanotechnol.* **7**, 51-55 (2012).
70 Kunding, A. H., Mortensen, M. W., Christensen, S. M. & Stamou, D. A fluorescence-based technique to construct size distributions from single-object measurements: application to the extrusion of lipid vesicles. *Biophys. J.* **95**, 1176-1188 (2008).
71 Lohr, C., Kunding, A. H., Bhatia, V. K. & Stamou, D. Constructing size distributions of liposomes from single-object fluorescence measurements. *Methods Enzymol* **465**, 143-160 (2009).
72 Ulbrich, M. H. & Isacoff, E. Y. Subunit counting in membrane-bound proteins. *Nat Methods* **4**, 319-321 (2007).
73 Heider, E. C., Peterson, E. M., Barhoum, M., Gericke, K. H. & Harris, J. M. Quantitative Fluorescence Microscopy To Determine Molecular Occupancy of Phospholipid Vesicles. *Analytical Chemistry* **83**, 5128-5136 (2011).
74 Uline, M. J. & Szleifer, I. Mode specific elastic constants for the gel, liquid-ordered, and liquid-disordered phases of DPPC/DOPC/cholesterol model lipid bilayers. *Faraday Discuss* **161**, 177-191; discussion 273-303 (2013).
75 van der Munnik, N. P., Sajib, M. S. J., Moss, M. A., Wei, T. & Uline, M. J. Determining the Potential of Mean Force for Amyloid-beta Dimerization: Combining Self-Consistent Field Theory with Molecular Dynamics Simulation. *J. Chem. Theory Comput.* **14**, 2696-2704 (2018).
76 Allender, D. W., Sodt, A. J. & Schick, M. Cholesterol-Dependent Bending Energy Is Important in Cholesterol Distribution of the Plasma Membrane. *Biophys. J.* **116**, 2356-2366 (2019).
77 Lorent, J. H. *et al.* Plasma membranes are asymmetric in lipid unsaturation, packing and protein shape. *Nat. Chem. Biol.* **16**, 644-652 (2020).
78 Szleifer, I. & Carignano, M. A. Tethered polymer layers. *Advances in Chemical Physics* (1996).
79 Szleifer, I., Kramer, D., Ben-Shaul, A., Gelbart, W. & Safran, S. A. Molecular theory of curvature elasticity in surfactant films. *J. Chem. Phys.* (1990).
80 Flory, P. J. *Statistical mechanics of chain molecules*. (Interscience Publishers, 1969).
81 Uline, M. J., Longo, G. S., Schick, M. & Szleifer, I. Calculating partition coefficients of chain anchors in liquid-ordered and liquid-disordered phases. *Biophys. J.* **98**, 1883-1892 (2010).
82 Grillo, D., Olvera de la Cruz, M. & Szleifer, I. Theoretical studies of the phase behavior of DPPC bilayers in the presence of macroions. *Soft Matter* **7** (2011).
83 Nap, R., Gong, P. & Szleifer, I. Weak polyelectrolytes tethered to surfaces: Effect of geometry, acid–base equilibrium and electrical permittivity. *Journal of polymer science Part B: Polymer Physics* **44**, 2638-2662 (2006).

### Supplementary Methods

#### Key principles in biosensor data extraction

Our high-content analysis allows us to extract the relative density of any biosensor at the plasma membrane of live cells in a spatially resolved manner. We image receptors and biosensors in 3D by XZY stacks, as we have previously employed this approach to accurately measure membrane topography and receptor density[20]. In the section below, key principles in extraction of biosensor signal at the plasma membrane are described in detail (Extended Data Figs. 1 and 2).

#### Classification of traces

For every XY-pixel in the XZY stack, we extract an intensity profile along the Z axis. This profile represents the sum of cytosolic and receptor-bound biosensor signal. In the case of zero receptor-bound biosensor signal, this profile is described by a sigmoid curve that is centred at the Z position of the plasma membrane ($Z_{PM}$). On the other hand, in the total absence of cytosolic signal all biosensors will be receptor-bound and give rise to a Gaussian-like profile. We leveraged this spatial phenotype of miniG and Nb80 to develop a physical model that encompasses the cytosolic and receptor-bound contributions (Extended Data Figs. 1a-c).

While a pure sigmoidal contribution is centred at $Z_{PM}$, increasing the contribution of receptor-bound component causes a left-shift of the intersect-at-half-maximum, or apparent Z-position ($Z_{app}$) (Extended Data Fig. 1c). The difference between $Z_{app}$ and $Z_{PM}$ is directly related to the contribution of the receptor-bound biosensors to the total intensity profile. Especially for traces where the Gaussian:Sigmoid contribution is small (i.e., the basal state, where most biosensor is cytosolic), $Z_{app} - Z_{PM}$ is a very sensitive readout for the presence of receptor-bound biosensors (Extended Data Fig. 1d). This sensitivity can be exploited up until the Gaussian:Sigmoid contribution is approximately 1:1. For Gaussian:Sigmoid contributions larger than 1, our sensitivity is not set by $Z_{app} - Z_{PM}$ (as shown by the plateau in Extended Data Fig. 1d), but rather by the increase in signal-to-noise of the Gaussian contribution.

As a negative control, we first measured $Z_{app} - Z_{PM}$ in the absence of β1AR for cells expressing only miniG$_s$ and the PM-label CellMask (Extended Data Fig. 1e). A comparison with the distribution $Z_{app} - Z_{PM}$ for cells expressing miniG$_s$ and β1AR readily shows a right shifted population, thus revealing the presence of XY-pixels that display miniG$_s$-bound receptors (c.f. Extended Data Figs. 1e and 1f). We used this negative control to determine a threshold ($Z_{thresh}$) for $Z_{app} - Z_{PM}$ measurements (dashed line in Extended Data Figs. 1e and 1f). All pixels above $Z_{thresh}$ reveal the presence of basal active conformations, whereas all pixels below $Z_{thresh}$ do not exhibit basal active conformations above our detection limit. The value of $Z_{thresh}$ results in a false-positive identification rate of 6%. This approach allowed us to classify and discriminate XY-pixels with and without a Gaussian contribution.

Next to the change in $Z_{app}$, we applied several additional criteria to classify intensity traces. Since cells contain unlabelled intracellular vesicles that result in strong signal depletions in the cytosolic fraction, we also classified these events. These classification criteria are summarised in Extended Data Table 1 and Extended Data Fig. 2.

Classification of traces primarily relies on the absence (class I and II, Extended Data Table 1) or presence (class III and IV, Extended Data Table 1) of receptor-bound biosensors, which is determined by $Z_{app}$ - $Z_{PM}$. To monitor the presence of intracellular vesicles, we introduced the following classification criteria. First, we calculated the slope of the last 15 pixels of the trace ($Slope_1$). Next, we defined $Slope_2$ as the slope of the trace between $Z_{PM}$ + 15 pixels (i.e., a diffraction limited step from $Z_{PM}$) and the start of $Slope_1$. $Slope_1$ detects the presence of intracellular vesicles at the end of the trace, whereas $Slope_2$ detects the presence of intracellular vesicles between $Z_{PM}$ and the end of the trace. Furthermore, we introduced a criterium that compares the intensity of the end of the trace ($I_{tail}$) with the average cytosolic intensity ($I_{cyt}$). $I_{cyt}$ is determined as the average signal of the cytosolic fraction of the biosensor signal. If $I_{tail}$ falls within the range of $I_{cyt} \pm \mathrm{sqrt}(I_{cyt})$, the tail of the trace is not subject to intracellular vesicles.

| No biosensor bound | | Receptor-bound biosensor | |
|---|---|---|---|
| **I** | **II** | **III** | **IV** |
| **Sigmoid** | **Sigmoid + cytosolic vesicle** | **Sigmoid + Gaussian** | **Sigmoid + Gaussian + cytosolic vesicle** |
| $Z_{app} - Z_{PM} < Z_{threshold}$ | $Z_{app} - Z_{PM} < Z_{threshold}$ | $Z_{app} - Z_{PM} > Z_{threshold}$ | $Z_{app} - Z_{PM} > Z_{threshold}$ |
| $Slope_1 < 2\%$ | $Slope_1 > 2\%$ | n.a. | $Slope_1 > 2\%$ |
| $Slope_2 < 2\%$ | $Slope_2 > 2\%$ | $Slope_2 < 10\%$ | $Slope_2 > 2\%$ |
| $I_{cyt} - \mathrm{sqrt}(I_{cyt}) < I_{tail}$ | n.a. | $I_{cyt} - \mathrm{sqrt}(I_{cyt}) < I_{tail}$ | n.a. |
| $I_{cyt} + \mathrm{sqrt}(I_{cyt}) > I_{tail}$ | $I_{cyt} + \mathrm{sqrt}(I_{cyt}) > I_{tail}$ | $I_{cyt} + \mathrm{sqrt}(I_{cyt}) > I_{tail}$ | $I_{cyt} + \mathrm{sqrt}(I_{cyt}) > I_{tail}$ |

**Extended Data Table 1 - Classification criteria for determining traces with and without receptor-bound biosensor.**

**Fitting of traces**

Following classification of the traces, all traces that have a Gaussian component (class III and IV, Extended Data Table 1) are fitted to extract the relative intensity of receptor-bound $miniG_s$ or Nb80 at the plasma membrane. We used Eq. 1 to fit all traces with 3 free parameters, i.e. A, B and D. We use the Z position of the plasma membrane for parameter C, we fixed the offset F as the average intensity of the first 5 pixels of the trace, and we described E in terms of B.

$$I(z) = \frac{A}{1+e^{-B(z-C)}} + \frac{De^{-0.25E^2(z-C)^2}}{1+E^2(z-C)^2} + F \qquad \text{(Eq. 1)}$$

In the presence of intracellular vesicles (class IV), we adapted Eq. 1 and introduced a corrective term that fits the depletion created by the vesicle, as shown in Eq. 2 below. This introduces 3 additional free parameters that are constrained between $Z_{PM}$ and the end of the trace. G is the amplitude of the corrective term, H the centre position of the vesicle, I and J are related to the width and shape of the vesicle. The I term can be described in terms of B.

$$I(z) = \frac{A}{1+e^{-B(z-C)}} + \frac{De^{-0.25E^2(z-C)^2}}{1+E^2(z-C)^2} + Ge^{-\frac{(z-H)^{2J}}{2I^{2J}}} + F \qquad \text{(Eq. 2)}$$

**Filtering of fits**

We have imposed several filtering criteria to select only high-quality fits, which allows us to extract the relative density of the biosensor with high accuracy. First, fits with an adjusted R-squared below 0.9 are removed from further analysis. Second, the standard error of the fit for the maximum intensity of the Gaussian must be smaller than 30% of the value of maximum intensity. The median standard error of the fit for the maximum intensity was 10% of the maximum intensity. Collectively, the above filtering typically accepts ~ 80 % of fits.

**Molecular mapping of membrane curvature-effects on GPCR density and allosteric activation**

As discussed extensively in reference [20], curvature has opposite effects on the inner/outer leaflets of the bilayer. This can be readily visualized on the 3D structure of the inactive β1AR where the hydrophobic contribution to receptor sorting density makes two marked stripes, positive/red on the outer leaflet and negative/blue on the inner Extended Data Fig. 15a. The receptor thus experiences a “tug-of-war” that for β1AR favors positive curvatures (Extended. Data Fig. 15b).

In comparison to sorting density, the curvature preference of the activation probability takes into account the energetic difference between active/inactive conformations as a function of curvature. In this scenario, the interleaflet tag-of-war is less important, and the outcome depends more on 1) what type of residues move between the two conformations, 2) how much they move, and 3) where they are placed along the lateral pressure profile (Extended Data Fig. 15c, d, e).

Our calculations revealed that the net increase in activation probability at negative mean curvature integrates contributions from residues located mostly in TM5 and TM6 (ED Fig.15c, f). Interestingly, many of them (e.g., K6.32, L6.34 and M6.43) have already been experimentally validated to increase spontaneous receptor activation through classical signaling and spectroscopic assays[22,39,67]. Previously, these constitutively active mutants were viewed through a receptor-centric perspective and interpreted to affect the intramolecular allosteric communication pathway between residues. Our results suggest a new mechanistic basis for these experimental results which comprises the energetic coupling between residues and membrane curvature.

**Chemicals for β$_2$AR liposome study**

1,2-dioleoyl-*sn*-glycero-3-phosphocholine (DOPC), 1,2-dioleoyl-*sn*-glycero-3-phospho-1′-*rac*-glycerol sodium salt (DOPG) and 1,2-dioleoyl-*sn*-glocero-3-phosphoethanolamine-N-biotinyl-(polyethylene glycerol-2000) ammonium salt (DSPE-PEG$_{2000}$-biotin were purchased from Avanti polar lipids. 5α-cholestan-3β-ol hemisuccinate (CHS) was purchased from Steraloids Inc. 1,1-dioctadecyl-3,3,3,3-tetramethylindodicarbocyanine perchlorate DiD-C18 oil (DiD) was purchased from Molecular probes. PLL(20)-*g*[3.5]-PEG(2) (PLL-PEG) and PLL(20)-*g*[3.5]-PEG(2)/PEG(3.4)-Biotin 18% (PLL-PEG-biot.) were purchased from (SuSoS, Switzerland). Isoproterenol (ISO) and Neutravidin were purchased from Sigma.

Buffer A: 100 mM NaCl, 20 mM HEPES pH 7.5

Buffer B: 1% octylglucoside (detergent),100 mM NaCl, 20 mM HEPES pH 7.5

**Preparation of β$_2$AR liposomes**

For liposomes used to study β2AR function, FLAG-tagged β2AR truncated after position 365 were expressed, purified, and labeled with BDY at Cys256 in the cytoplasmic end of TM6, as described earlier for other fluorophores[68]. Briefly, liposomes containing DOPC/CHS/DOPG/DiD/DSPE-PEG$_{2000}$-Biotin (79.5:10:10:0.5:0.1) were prepared by evaporating chloroform under argon, and subsequently dried for 1 hour in vacuum to obtain a thin lipid film. The film was resuspended in buffer A and liposomes were formed during sonication for 1 hour in an ice water bath. Labeled β$_2$AR-BDY in buffer B and liposomes (1 g/l) were mixed to a 1:1000 protein to lipid ratio in sample buffer B until 300 μl and kept on ice for 2 hours. Liposomes were formed by the removal of detergent on a Sephadex G-50 column (25x0.8 cm).

**Surface functionalization for the β2AR liposome study**

Surfaces used for the β2AR function study were prepared as described previously[69-71]. Ultra clean glass slides (thickness 170 +/-10 μm) were plasma etched for 2 min., mounted in microscope chamber and incubated for 30 min. with a mixture of 1 g/l PLL-PEG and PLL-PEG-Biotin (dissolved n 15 mM HEPES at pH 5.6) in a ratio of 1000:6 for β$_2$AR surfaces. Unbound polymers were washed away with buffer A and the surfaces were incubated with 0.1 g/l Neutravidin (dissolved in 15 mM HEPES) for 10 min. followed by extensively washing with buffer A. The density of immobilized liposomes was controlled by recording liposome binding in real time after adding liposomes 4 µl 0.05 g/l to a 80 µl chamber. Unbound liposomes were removed by washing when the surface density reached approximately 500 liposomes per 81.76×81.76 µm$^2$.

**Size calibration of β2AR liposomes**

Diameters of imaged fluorescent liposomes were determined as described previously[70]. In brief, the integrated intensity of a single liposome ($I_M$) is proportional to the number of fluorescently labeled lipids in the liposome membrane, and thereby proportional to the liposome surface area ($A_{ves}$). The diameter ($D$) is consequently related to ($I_M$) by the proportionality factor ($C_{cal}$) according to:

$$I_M \propto A_{ves} = \pi D^2 \quad \Rightarrow \quad D = C_{cal}\sqrt{I_M}$$

$C_{cal}$ was quantified with a calibration sample where the liposomes were extruded 20 times through two 50 nm polycarbonate filters (Millipore) that produced a narrow size distribution. The calibration sample was first examined by dynamic light scattering (DLS, ALV-5000 Correlator equipped with a 633 nm laser line) followed by TIRF microscopy using identical membrane imaging conditions as for a normal sample. The mean integrated membrane intensity of the imaged calibration sample was correlated to the mean radius obtained by DLS. Once the calibration factor was known all integrated liposome intensities were converted into physical liposome diameter (SI Fig. 1e).

To convert liposome diameter ($D$) into membrane curvature (MC) of the liposome, we first changed the unit from nm to µm and calculated the membrane curvatures and the propagated error of these according to:

$$MC = \frac{1}{R} = \frac{1}{\frac{1}{2}D} = \frac{2}{D}$$

$$\delta MC = \sqrt{\left(\frac{\partial MC}{\partial D}\delta D\right)^2} = \sqrt{\left(-\frac{2}{D^2}\delta D\right)^2}$$

**TIRF microscopy on β2AR liposomes**

All images of receptor-liposome complexes as well as BDY bleaching were acquired on a Leica DMI6000 TIRF system using an oil immersion objective HCX PL APO CS with x100 magnification and numerical aperture of 1.46. Membrane label DiD and receptor label BDY were excited with a 633 nm laserline, and a 488 nm laserline respectively. BDY emission was selected with a dichroic mirror Q495LP and a bandpass filter HQ525/50m. DiD emission was selected with a dichroic mirror Q660LP and a bandpass filter HQ700/75m. All filters and dichroics were purchased from Chroma technology, Brattleboro, VT. Fluorescence intensities were collected on an electron-multiplying Andor Ixon 897 camera. Images were acquired at a resolution of 512×512 pixels each pixel corresponding to 160×160 nm at a bit-depth of 14. Membrane intensities were imaged using an exposure time of 400 ms. Bleaching traces of BDY were acquired using an exposure time of 250 ms, each frame was acquired and transferred in 304 ms. Static images of BDY receptor function were acquired using 400 ms in exposure time.

**Single step bleaching analysis of β2AR liposomes and image processing**

We calibrated receptor densities using single molecule bleaching traces of BDY according to [72,73]. Bleaching traces of single liposomes were acquired using a TIRF microscope setup (see description under TIRF microscopy). Single molecule bleaching trace intensities were extracted using ROI integration of BDY with software written in Igor Pro Ver. 6.01 (Wavemetrics, Tigard, OR). The single molecule intensity step was quantified by subtracting its average step intensity by its average background intensity. To correlate the obtained step amplitude to intensities extracted in the receptor activation study, the same software also extracted membrane and BDY intensities for each receptor-liposome complex using 2D Gaussian integration from images acquired under conditions used for the receptor function study. A correlation between 2D Gaussian integrated intensities and ROI integrated intensities was found and the linear intensity regime of BDY provided a factor we used to convert the obtained single molecule step using ROI integration into single steps of 2D Gaussian integration. We found that the collected bleaching steps to be Gaussian distributed, suggesting identical photophysics of the fluorophores of the bleaching steps. This allowed us to quantify the number of receptors for each liposome by dividing the 2D Gaussian intensity of each liposome with the mean single step amplitude (SI Fig. 1f).

**Image processing of receptor activity of $\beta_2$AR in liposomes**

For quantifying the receptor orientation as a function of liposome for each surface immobilized liposome-receptor complex, a pair of 2 images (DiD, BDY) were collected at each microscope chamber position before and incubating the sample with KI quencher (20 mM). All reported intensities with errors in these studies were extracted from the fluorescence images using the software (described above) written in Igor Pro Ver. 6.01 (Wavemetrics, Tigard, OR). For each image, the diffraction limited liposome intensity spots above a user-defined threshold were identified and fitted with a 2D Gaussian in order to integrate the intensities and assign $(x_0,y_0)$

coordinates to each spot center. To colocalize the related membrane and receptor intensity spots between the 4 images, the center positions were matched and particles were accepted if two consecutive centers were within a certain defined pixel distance. Only liposomes colocalized in all four images were included in the analysis (SI Fig. 1c and d).

A 2D Gaussian fit to each liposome intensity distribution provided liposome position ($x_0$,$y_0$), amplitude (A), widths ($\sigma_x$, $\sigma_y$) and correlation in x and y direction ($\rho$) with standard errors for the individual parameters according to:

$$G(x,y) = z_0 + A\exp\left[\frac{-1}{2\left(1-\rho^2\right)}\left(\left(\frac{x-x_0}{\sigma_x}\right)^2 + \left(\frac{y-y_0}{\sigma_y}\right)^2 - \left(\frac{2\rho(x-x_0)(y-y_0)}{\sigma_x\sigma_y}\right)\right)\right]$$

The circularity of the liposome was calculated as the ratio of the major and minor axis of the intensity distribution independent of liposome elongation orientation. The intensity $I$ and its standard error $\delta I$ of the fluorescence intensity distribution was calculated from fit parameters according to:

$$I = 2\pi A\sigma_x\sigma_y\sqrt{1-\rho^2}, \quad \delta I = \sqrt{\left(\frac{I}{A}\delta A\right)^2 + \left(\frac{I}{\sigma_x}\delta\sigma_x\right)^2 + \left(\frac{I}{\sigma_y}\delta\sigma_y\right)^2 + \left(\frac{-I\rho}{1-\rho^2}\delta\rho\right)^2}$$

To quantify the receptor activation (increase of BDY intensity) as a function of liposome diameter, two imaging methods were employed.

1. IM. First imaging method included imaging of membrane and receptor intensities at different positions in the microscope chamber before and after incubation with ISO.
2. IM. Second imaging method included a set of 3 images that was acquired using a fixed position of the image frame: membrane, BDY before and after incubation with ISO.

**Molecular-field Theory (MFT)**

To elucidate molecular mechanisms for plasma membrane curvature modulation of the probability of basal activation of a β1AR protein, we use a highly detailed MFT to determine the physical properties of curved asymmetric lipid bilayers with a β1AR protein embedded within its structure. Recently, the MFT was successfully utilized to capture how plasma membrane mean curvature couples entropic and enthalpic interactions to drive the inactive structure of the β1AR protein to laterally organize (spatial density inhomogeneities) on the surface of living cells[20] in quantitative agreement with experiments. Previous versions of the

MFT were shown to be in excellent agreement with experiments on N-Ras anchor partitioning into liquid-ordered (lo) versus liquid-disordered (ld) phases on liposomes as a function of curvature[42]. The lipid bilayers were comprised of three components, PSM, DOPC, and cholesterol, and the quantitative comparisons were very strong considering there is only one fitting parameter in the MFT. Another two MFT and experimental studies on curvature sensing that produced similar levels of quantitative agreement were on N-Ras anchors binding to pure component liposomes comprised of DLPC, DMPC, POPC, and DOPC in the ld phase[43], and N-Ras, Synaptotagmin-1, and Annexin-12 binding to pure DOPC bilayers[44]. In all these studies, the MFT demonstrated that the lateral pressure profile in the lipid bilayer could be used to make accurate predictions on the curvature sensing of proteins with a variety of binding domains in several diverse lipid environments.

The MFT uses a free energy functional that is constructed by explicitly writing each of the energetic/entropic contributions and then minimizing the free energy with respect to the free variables. There is only one fitting parameter used in the calculation, and that is the strength of the hydrophobic interactions between $CH_2$ and $CH_3$ groups of the lipids or proteins. We input the physical conformations of the chains with the conformations of the protein and through free energy minimization we obtain the probability of each of those conformations as a function of the constraints imposed on the system. Through this method we can obtain the molecular level equilibrium physical parameters that we need to elucidate the fundamental molecular driving forces for protein localization and basal activation.

The basic concept of the theory is to consider each possible conformation of the lipids around the β1AR protein and formulate a free energy in terms of the probability of each of those conformations. By summing over each possible conformation, we are explicitly including fluctuations into the calculation. The intramolecular interactions are therefore treated exactly within the model. The intermolecular interactions are only exact within the length scale of a single molecule, so correlations beyond that length scale are only approximate. We are using a field theory that includes the physical conformations of the molecules and fluctuations, and we expect the agreement that we see with the experiments to be due to these improvements over more simplified mean-field theories[20,42-44,74,75].

**Theoretical calculations of our model of the plasma membrane as a function of curvature.**

The basic concept of the theory is to consider each possible conformation of the lipids and formulate a free energy in terms of the probability of each of those conformations. The minimal free energy provides us with an explicit expression for the probability of each molecular conformation in terms of the intermolecular and intramolecular interactions. The chemical potential of each lipid species in each leaflet is constrained to be constant for each curvature. The chemical potential of cholesterol is set equal across the leaflets so that in this model, cholesterol is the only molecule that can flip between the two leaflets. The inner leaflet (cytosolic leaflet) has the lipid chemical potentials set so that for a flat membrane the mol fraction of DOPS is 0.35, the mole fraction of DOPE is 0.20, POPC is 0.20, PiP2 is 0.02, and cholesterol is at 0.23. For the outer leaflet (non-cytosolic leaflet) the lipid chemical potentials are set so that for a flat membrane the mol fraction of POPC is 0.33, the mol fraction of PSM is 0.33, and the mol fraction of cholesterol is 0.33. These compositions are selected from recent experiment, theory and simulation work on the plasma membrane[76,77]. The conformations of the chains are described by Flory's Rotational Isomeric States model[78-80] in which each $CH_2$ group is in one of three configurations: the lowest energy *trans*, or the *gauche-plus* or *gauche-minus*, both of which are of an energy, 500 cal/mol, greater than that of the *trans* configuration. The plasma membrane is in contact with a salt solution of *NaCl* in water. The water is treated as dissociable from the neutral $H_2O$ into $H^+_{(aq)}$ and $OH^-_{(aq)}$, each of the three with a volume of 0.03 nm$^3$. It is convenient to define a local volume fraction, $\phi_i(z)$, which is a function of position, $z$, and is related to the local number density, $\rho_i(z)$, by $\phi_i(z) = \rho_i(z)v_i$, where $v_i$ is the molecular volume of that chemical species.

We define the *z*-axis to be perpendicular to the interfaces with its origin at the midplane of the bilayer. The total area at distance *z*, $A(z,\mathbf{c})$, parallel to the midplane of the bilayer, is given by[44,81]:

$$A(z,\mathbf{c}) = A(0)[1 + (c_1 + c_2)z + c_1c_2z^2] \qquad \text{(S1)}$$

where $c_1$ and $c_2$ are the principal curvatures, $c_i = 1/r_i$, and $r_i$ is the radius of curvature of the midplane of the bilayer. The intermolecular repulsions are modelled as excluded volume interactions and therefore we include them through packing constraints. Namely, we solve our molecular theory under the constraint that the total density is constant at each position *z* (i.e., that it is incompressible)[78-83]. The constant density constraint implies that the average volume fraction of the molecules in any layer must be equal to unity. That is,

$$\sum_{\delta} \langle \phi_{L,\delta}(z) \rangle + \phi_w(z) + \phi_{H^+}(z) + \phi_{OH^-}(z) + \phi_+(z) + \phi_-(z) = 1 \qquad \text{(S2)}$$

where $\delta = I,E$ $\delta = I,E$ $\delta = I,E$ $\delta = I,E$ and *I* and *E* stand for the interior and exterior leaflet of the bilayer, respectively. The subscript (+) stands for the dissociated sodium cation and the subscript (–) stands for the dissociated chlorine anion. Throughout the rest of this section, we will use $x_{L,\delta}$ to denote the mole fraction of lipids in leaflet δ.

The molecular theory uses a free energy functional that is constructed by explicitly writing each of the energetic/entropic contributions and then minimizing the free energy with respect to the free variables (for more details please see previous work with this model[20,78-83]). We input the physical conformations of the chains and through free energy minimization we obtain the probability of each of those conformations as a function of the constraints imposed on the system. Through this method we can obtain the molecular level equilibrium physical parameters that we need to elucidate the fundamental molecular driving forces for protein localization. The free energy of a membrane is given by:

$$
\begin{aligned}
\frac{\beta W}{N_L} &= \sum_{\delta} x_{L\delta}\left[\ln\left(\frac{x_{L\delta}\lambda_L^2}{a(0)}\right)-1\right] + \sum_{\delta}\sum_{\alpha_\delta} 2x_{L\delta}\left[P_{\delta,t}(\alpha_\delta)\left(\beta\varepsilon_t(\alpha_\delta)+\ln\left(P_{\delta,t}(\alpha_\delta)\right)\right)\right] \\
&+\sum_{\delta}\sum_{\sigma_\delta} x_{L\delta}\left[P_{\delta,h}(\sigma_\delta)\left(\ln\left(P_{\delta,h}(\sigma_\delta)\right)\right)\right] + \frac{\beta}{N_L}\int\left[\langle\rho_q(\mathbf{r})\rangle\psi(\mathbf{r}) - \frac{\varepsilon}{2}(\nabla\psi(\mathbf{r}))^2\right]d\mathbf{r} \\
&+\sum_{\delta}\sum_{\delta'}\frac{\beta\chi}{2N_L}v_{L,t}\int\langle\rho_{L\delta,t}(\mathbf{r})\rangle\langle\rho_{L\delta',t}(\mathbf{r})\rangle d\mathbf{r} + \frac{\beta\chi}{2N_L}v_{L,t}\int\langle\rho_{L,I,h,c}(\mathbf{r})\rangle\langle\rho_{L,I,h,c}(\mathbf{r})\rangle d\mathbf{r} \\
&+\frac{\beta\chi}{2N_L}v_{L,t}\int\langle\rho_{L,E,h,c}(\mathbf{r})\rangle\langle\rho_{L,E,h,c}(\mathbf{r})\rangle d\mathbf{r} + \frac{1}{N_L}\int\rho_+(\mathbf{r})\left[\ln\left(\rho_+(\mathbf{r})v_w\right)-1-\beta\mu_+\right]d\mathbf{r} \\
&+\frac{1}{N_L}\int\rho_-(\mathbf{r})\left[\ln\left(\rho_-(\mathbf{r})v_w\right)-1-\beta\mu_-\right]d\mathbf{r} + \frac{1}{N_L}\int\rho_{H^+}(\mathbf{r})\left[\ln\left(\rho_{H^+}(\mathbf{r})v_w\right)-1\right]d\mathbf{r} \\
&+\frac{1}{N_L}\int\rho_{OH^-}(\mathbf{r})\left[\ln\left(\rho_{OH^-}(\mathbf{r})v_w\right)-1\right]d\mathbf{r} + \frac{1}{N_L}\int\rho_w(\mathbf{r})\left[\ln\left(\rho_w(\mathbf{r})v_w\right)-1\right]d\mathbf{r} + \\
&\frac{1}{N_L}\int\beta\pi(\mathbf{r})\left[\sum_{\delta}\langle\phi_{L\delta}(\mathbf{r})\rangle + \phi_w(\mathbf{r}) + \phi_{H^+}(\mathbf{r}) + \phi_{OH^-}(\mathbf{r}) + \phi_+(\mathbf{r}) + \phi_-(\mathbf{r}) - 1\right]d\mathbf{r}
\end{aligned}
$$

(S3)

where $W$ is the constrained free energy of the system, $P_{\delta,t}(\alpha_\delta)$ is the probability of the fatty-acid tails in leaflet δ, $N_L$ is the total number of lipids, $\alpha_\delta$ is the conformation of the fatty-acid chain of the lipid molecule, $\sigma_\delta$ is the conformation of the lipid headgroups, $\varepsilon_t$ is the internal energy of the chain that arises from having *gauche* dihedral angles in a particular conformation, $v_{L,t}$ is the volume of a $CH_2$ unit that is equal to 0.027 nm$^3$, and $\beta = 1/k_B T$. The first term in the above equation is the contribution to the total free energy from the translation of the lipids, where $\lambda_L$ is the thermal de Broglie wavelength of the lipid molecules. The second term accounts for the conformational entropy and internal energy of the chains. The third term accounts for the conformational entropy of the headgroups. We modelled the PC headgroup of the lipids in a very similar way to previous publications using this theory[20,74], where instead of modelling the volume of the headgroup as two spheres we again used the Rotational Isomeric States model (RIS). The two glycerol carbons bonded to the phosphate group are fixed in space, and the remainder of the headgroup, comprised of two $CH_2$ groups, and a tri-methylated tertiary amine group, move according to the RIS model. The location of the phosphate group has a fixed charge of -1*e*, and the movable amine group has a fixed charge of +1*e*, where *e* is the fundamental charge. The PE headgroups were modelled very similarly to the PC groups except for the 3 $CH_3$ groups attached to the amine group are replaced by 3 protons that greatly reduce both the volume and hydrophobicity of the headgroup. The PS headgroup takes the PE headgroup and adds a hydrogen and a carboxylic acid group to the carbon bonded to the amine group. The acid group adds an additional negative charge to the headgroup at pH 7. The fourth term is the contribution of electrostatics[20,44,83], where $\psi(\mathbf{r})$ is the electrostatic potential, and $\varepsilon$ is the dielectric coefficient, which is taken to be that of water (78.5 times the value of the dielectric value of vacuum). $\langle \rho_q(\mathbf{r}) \rangle$ is the ensemble averaged number density of charges given by:

$$\langle \rho_q(\mathbf{r}) \rangle = \sum_{\delta} \langle \rho_{L\delta,h,q}(\mathbf{r}) \rangle e + \rho_+(\mathbf{r})e + \rho_{H^+}(\mathbf{r})e - \rho_-(\mathbf{r})e - \rho_{OH^-}(\mathbf{r})e, \qquad \text{(S4)}$$

where $\langle \rho_{L\delta,h,q}(\mathbf{r}) \rangle$ is the local density of charges on the headgroups. The fifth, sixth, and seventh terms are the contributions of the hydrophobic interactions in the lipids system. $\langle \rho_{L\delta,h,c}(\mathbf{r}) \rangle$ is the local density of $CH_2$ and $CH_3$ groups in the headgroups of the lipids. $\chi = -5k_BT$ is the strength of the hydrophobic interactions. The eighth and ninth terms in the free energy represent the translational free energy of the cations and anions of the salt as well as the bulk chemical potential of both species (the chemical potential is used to impose a bulk salt concentration of 0.1M). The tenth, eleventh, and twelfth terms represent the translational free energy of the dissociated $H^+_{(aq)}$ and $OH^-_{(aq)}$ species along with that of the water. The bulk pH in all the calculations is set at 7 and the pK$w$ is set at 14.0. The final term accounts for the repulsive steric interactions that are accounted for in an incompressibility constraint. $\pi(\mathbf{r})$ is a Lagrange multiplier that is introduced to enforce the incompressibility constraint.

We will now use the coordinate $z$ in the place of $\boldsymbol{r}$ to simplify the notation. When we use $z$ for the remainder of the text, we mean the full 3-dimensional space. The probability distributions for the headgroups and fatty-acid tails of the lipids (here written with the explicit curvature dependence) are:

$$P_{\delta,t}(\alpha_\delta,\mathbf{c}) = \frac{1}{q_{\delta,t}(\mathbf{c})}\exp\left[\begin{array}{l} -\beta\varepsilon_t(\alpha_\delta) - \int \beta\pi(z,\mathbf{c})v_{L\delta,t}(z,\alpha_\delta)dz - \\ \beta\chi v_{L,t}\int\left[\langle \rho_{L,I,t}(z,\mathbf{c})\rangle + \langle \rho_{L,E,t}(z,\mathbf{c})\rangle\right] n_{t,\delta}(z,\alpha_\delta)dz \end{array}\right]$$

$$P_{\delta,h}(\sigma_\delta,\mathbf{c}) = \frac{1}{q_{\delta,h}(\mathbf{c})}\exp\left[\begin{array}{l} -\int \beta\pi(z,\mathbf{c})v_{L\delta,h}(z,\sigma_\delta)dz - \int \beta\psi(z,\mathbf{c})en_{L\delta,h}(z,\sigma_\delta)dz \\ -\beta\chi v_{L,t}\int\left[\langle \rho_{L\delta,h}(z,\mathbf{c})\rangle\right] n_{h,\delta}(z,\sigma_\delta)dz \end{array}\right]$$

(S5)

where $q_{\delta,t}(\mathbf{c})$ and $q_{\delta,h}(\mathbf{c})$ are the single molecule partition functions (normalization of the probability) defined as

$$q_{\delta,t}(\mathbf{c}) = \sum_{\alpha_\delta} \exp\left[\begin{array}{l} -\beta\varepsilon_t(\alpha_\delta) - \int \beta\pi(z,\mathbf{c}) v_{L\delta,t}(z,\alpha_\delta)dz - \\ \beta\chi v_{L,t} \int \left[\langle \rho_{L,I,t}(z,\mathbf{c})\rangle + \langle \rho_{L,E,t}(z,\mathbf{c})\rangle\right] n_{t,\delta}(z,\alpha_\delta)dz \end{array}\right]$$

$$q_{\delta,h}(\mathbf{c}) = \sum_{\sigma_\delta} \exp\left[\begin{array}{l} -\int \beta\pi(z,\mathbf{c}) v_{L\delta,h}(z,\sigma_\delta)dz - \int \beta\psi(z,\mathbf{c}) e n_{L\delta,h}(z,\sigma_\delta)dz \\ -\beta\chi v_{L,t} \int \left[\langle \rho_{L\delta,h}(z,\mathbf{c})\rangle\right] n_{h,\delta}(z,\sigma_\delta)dz \end{array}\right] \qquad \text{(S6)}$$

$\beta\pi(z,\mathbf{c})$ and $\beta\psi(z,\mathbf{c})$ are determined self-consistently for each curved state of the bilayer, and these field variables explicitly control the probability distributions of the lipid tails and headgroups.

**Theoretical calculations of the basal activation of β1AR as a function of curvature.**

This is a two-state model for a receptor that exists in an inactivated state and an activated state signified by superscript, *. The probability of the inactive state is $P(\vec{c})$. The probability of the active state is $P^*(\vec{c})$, and both states are an explicit function of curvature, $\vec{c}$. $I(\vec{c})$ is the interaction energy of the inactive state protein with all the lipid and solvent molecules in a given state of curvature, and $I^*(\vec{c})$ is the interaction energy of the active state protein with all of the lipid and solvent molecules in a given state of curvature as described above.

Assuming that the number of proteins, $N_P$, is constant and that the concentration of proteins is in infinite dilution, then we can write the free energy as

$$\frac{F(\vec{c})}{N_P k_B T} = P(\vec{c})[lnP(\vec{c}) + \beta I(\vec{c})] + P^*(\vec{c})[lnP^*(\vec{c}) + \beta I^*(\vec{c}) + \beta\varepsilon^*] + \alpha[P(\vec{c}) + P^*(\vec{c}) - 1],$$

where $\varepsilon^*$ is the intramolecular energy difference of the protein in the active state relative to the inactive state, and $\alpha$ is a Lagrange multiplier enforcing normalization of the probability.

We can minimize the free energy to obtain the probabilities,

$$P(\vec{c}) = \frac{\exp(-\beta I(\vec{c}))}{\exp(-\beta I(\vec{c})) + \exp(-\beta\varepsilon^* - \beta I^*(\vec{c}))}\ ;\ P^*(\vec{c}) = \frac{\exp(-\beta\varepsilon^* - \beta I^*(\vec{c}))}{\exp(-\beta I(\vec{c})) + \exp(-\beta\varepsilon^* - \beta I^*(\vec{c}))}\ .$$

The relative probability of activation as a function of curvature is,

$$\frac{P^*(\vec{c})}{P(\vec{c})} = \exp\{-\beta[\varepsilon^* + I^*(\vec{c}) - I(\vec{c})]\}\ .$$

The local interaction energies, $I(\vec{c})$ or $I^*(\vec{c})$, of β1AR can be calculated at infinite dilution from the molecular theory where the hydrophobic distribution, shape, charge distribution, and total volume the protein is explicitly considered. An assumption is made that the total chemical potential of β1AR is enforced at every point in the plasma membrane to be equal to the value when the membrane is flat. This assumption allows one to calculate the local protein concentration as a function of curvature by,

$$I(\vec{c}) = -\int \beta\pi(z,\vec{c})v_p(z)dz - \int \beta\psi(z,\vec{c})en_{p,q}(z)dz - \beta\chi v_{L,t}\int\langle\rho_{L,\delta,h}(z,\vec{c})\rangle n_{p,c}(z)dz, \quad \text{(S7)}$$

where $v_p(z)dz$ is the volume contribution of the protein at position $z$ in the bilayer. $n_{p,c}(z)dz$ is the number of $CH_2$ and $CH_3$ groups in the protein located at position $z$, and $n_{p,q}(z)dz$ is the number of charged units in the protein located at position $z$. The integration over $v_p(z)dz$ from $z_{min}$ to $z_{max}$ yields the total volume of the protein. From this expression, it is evident that the size, shape, hydrophobicity, and charge distributions are critical parameters for determining the relative interactions surrounding the inactive and active states of the protein.

The structure of inactive β1AR was produced using PDB structure file (PDB ID: 2YCW). The structure of the active β1AR was produced using PDB structure file (PDB ID: 6H7J). There exist several residue numbers between the two structures that do not agree, and when that occurs, we use the 2YCW residue identity for that number. The positions of the atoms as a function of distance perpendicular to the *xy*-plane of the bilayer headgroups are then exactly determined. The volumes of the individual units are calculated by using the approximate van der Waals radii of 0.16 nm for carbon, 0.14 nm for oxygen, 0.145 nm for nitrogen, 0.173 nm for sulfur, and 0.05 nm for hydrogen. The locations of $CH_2$ and $CH_3$ groups contributed to $n_{p,c}(z)$ and the locations of the charged amino acids contributed to $n_{p,q}(z)$.
